# Functional Landscape of SARS-CoV-2 Cellular Restriction

**DOI:** 10.1101/2020.09.29.319566

**Authors:** Laura Martin-Sancho, Mary K. Lewinski, Lars Pache, Charlotte A. Stoneham, Xin Yin, Dexter Pratt, Christopher Churas, Sara B. Rosenthal, Sophie Liu, Paul D. De Jesus, Alan M. O’Neill, Anshu P. Gounder, Courtney Nguyen, Yuan Pu, Aaron L. Oom, Lisa Miorin, Ariel Rodriguez-Frandsen, Matthew Urbanowski, Megan L. Shaw, Max W. Chang, Christopher Benner, Matthew B. Frieman, Adolfo García-Sastre, Trey Ideker, Judd F. Hultquist, John Guatelli, Sumit K Chanda

## Abstract

A deficient interferon response to SARS-CoV-2 infection has been implicated as a determinant of severe COVID-19. To identify the molecular effectors that govern interferon control of SARS-CoV-2 infection, we conducted a large-scale gain-of-function analysis that evaluated the impact of human interferon stimulated genes (ISGs) on viral replication. A limited subset of ISGs were found to control viral infection, including endosomal factors that inhibited viral entry, nucleic acid binding proteins that suppressed viral RNA synthesis, and a highly enriched cluster of ER and Golgi-resident ISGs that inhibited viral translation and egress. These included the type II integral membrane protein BST2/tetherin, which was found to impede viral release, and is targeted for immune evasion by SARS-CoV-2 Orf7a protein. Overall, these data define the molecular basis of early innate immune control of viral infection, which will facilitate the understanding of host determinants that impact disease severity and offer potential therapeutic strategies for COVID-19.

## INTRODUCTION

The ongoing coronavirus disease 2019 (COVID-19) pandemic, caused by severe acute respiratory syndrome coronavirus 2 (SARS-CoV-2), is responsible for a reported 23.5 million infections, and over 800,000 deaths worldwide as of this writing (Dong et al., 2020). Following infection with SARS-CoV-2, COVID-19 clinical presentation ranges from asymptomatic or mild (suggested to account for ∼ 80% of infections), to severe disease that typically requires hospitalization and assisted respiration (Huang et al., 2020). While age and co-morbidities, such as obesity and cardiovascular disease, have been linked to COVID-19 severity, recent data suggest that cellular immune responses to viral infection are also a critical determinant of disease outcome (Mathew et al., 2020). For instance, loss-of-function mutations in the immune sensor *TLR7* and downregulation of the type I interferon (IFN) response have been associated with severe COVID-19 (van der Made et al., 2020). In addition, two recent studies that conducted an integrated immune analysis of COVID-19 patients found impaired IFN responses in severe and critically ill patients (Arunachalam et al., 2020; Hadjadj et al., 2020). Further support for the role of IFN in COVID-19 outcome comes from a study of 127 patients receiving interferon beta-1b in combination with lopinavir–ritonavir and ribavirin, which reported lower SARS-CoV-2 viral load and shedding in the lungs and reduced length of hospitalization (Hung et al., 2020). Taken together, these data underscore an emerging role for IFN-mediated cellular responses in the control of SARS-CoV-2 infection and COVID-19 severity.

Viral infection is sensed by pattern-recognition receptors (PRR), which initiate a signaling cascade that produces cytokines, including IFN. Binding of IFN to its receptor (IFNAR) promotes the transcriptional activation of hundreds of interferon stimulated genes (ISGs), many of which exert antiviral activities (Schoggins et al., 2011). Concerted expression and regulation of these PRRs and downstream signaling molecules, transcription factors, and effectors are necessary to mount a successful antiviral response. Thus, viruses have developed various strategies to interfere with and evade these antiviral programs (García-Sastre, 2017). Recent work has shown that SARS-CoV-2 infection is sensitive to IFN treatment, as RNAseq of COVID-19 patients samples and *in vitro* infection models revealed upregulation of ISGs (Blanco-Melo et al., 2020; Emanuel et al., 2020; Lamers et al., 2020; Overmyer et al., 2020; Sun et al., 2020). In addition, the ISG *LY6E* has been identified as a negative regulator of SARS-CoV-2 (Pfaender et al., 2020), and the ISGs *AXIN2, CH25H, EPSTI1, GBP5, IFIH1, IFITM2*, and *IFITM3* were found to block entry of a pseudotyped vesicular stomatitis virus (VSV) harboring SARS-CoV-2 Spike (S) protein (Zang et al., 2020). Ultimately, a comprehensive evaluation of ISGs that inhibit infection of SARS-CoV-2 will be necessary to understand the cellular control of viral infection and their potential impact on COVID-19 outcome.

To uncover the cellular antiviral response to SARS-CoV-2 infection, we conducted a gain-of-function screen using 399 human ISGs. These data revealed that restriction of SARS-CoV-2 is mediated by a limited subset of 65 ISGs, most of which reside in the ER or Golgi compartments and function to regulate endoplasmic reticulum-associated protein degradation (ERAD), lipid membrane composition, and vesicle transport. Among these was BST2, found to inhibit viral egress and to be antagonized by SARS-CoV-2 accessory protein Orf7a to rescue virion release. The identification of the ISG subset that direct the antiviral activity of IFN illuminates the molecular and genetic determinants of early immune regulation that contribute to COVID-19 outcome, and provide attractive specific targets for therapeutic intervention.

## RESULTS AND DISCUSSION

### IFN-mediated restriction of SARS-CoV-2 relies on a limited subset of ISGs

To define the cellular effectors that act to limit SARS-CoV-2 infection, we first sought to determine which genes are activated upon IFN stimulation (hereafter referred to as ISGs) in disease-relevant cell types. Human tracheobronchial epithelial (HTBE) and human alveolar epithelial A549 cells were treated with IFN for 8 h and then subjected to RNAseq. Using cut-off criteria of log2FC>1.5 and p value < 0.05, we identified 139 ISGs upregulated in HTBE, 121 ISGs upregulated in A549 cells, and 152 ISGs upregulated in both HTBE and A549 (**Fig S1A**). This dataset encompassed ISGs with previously characterized broad-acting antiviral activities that included *MX1, OAS1, OASL* and *IFI6* (Hubel et al., 2019). In addition, Schoggins *et al*. previously assembled a list of 387 curated ISGs, of which 149 overlapped with the HTBE/A549 dataset (**Fig S1B**) (Schoggins et al., 2011). We combined these experimental and published datasets, and identified 399 ISGs as available, validated, and full-sequence length cDNA clones (**Fig S1B, Table S1A, B**).

Next, we evaluated the ability of these 399 ISGs to inhibit SARS-CoV-2 replication using ectopic expression screening. These studies were conducted using the human epithelial cell line 293T, as these cells can be transfected with high efficiencies and support productive replication of SARS-CoV-2 when expressing the viral entry factors *ACE2* and *TMPRSS2* (Hoffmann et al., 2020). 293T cells were transfected with individual ISGs along with *ACE2* and *TMPRSS2* for 30 h, and then challenged with SARS-CoV-2 at a low multiplicity of infection (MOI = 0.0625). Cells were fixed at 40 h post-infection, and infectivity was determined using immunostaining for SARS-CoV-2 nucleoprotein (N) (**Fig 1A**). cDNA encoding chloramphenicol acetyltransferase (*CAT*) was included on each plate as negative control, and cDNA encoding the SARS-CoV-2 negative regulator *LY6E* (Pfaender et al., 2020) was included as positive control (**Fig 1A, B**). Screens were conducted in duplicate and showed good reproducibility with a Pearson correlation coefficient (r) = 0.81 (**Fig 1C**). After applying cut-off criteria for infectivity (log2FC at least four standard deviations lower than the *CAT* negative control) and cell viability (at least 70% number of cells of the negative control), we identified 65 ISGs that inhibited SARS-CoV-2 replication (**Fig 1B**). Cross-comparison of these 65 factors with published datasets of upregulated genes from COVID-19 patient samples and *in vitro* infected lung cell models revealed a significant overlap (**Fig S1C**), suggesting that these factors are also stimulated in response to SARS-CoV-2 infection (Blanco-Melo et al., 2020; Emanuel et al., 2020; Overmyer et al., 2020; Sun et al., 2020).

**Figure 1.**
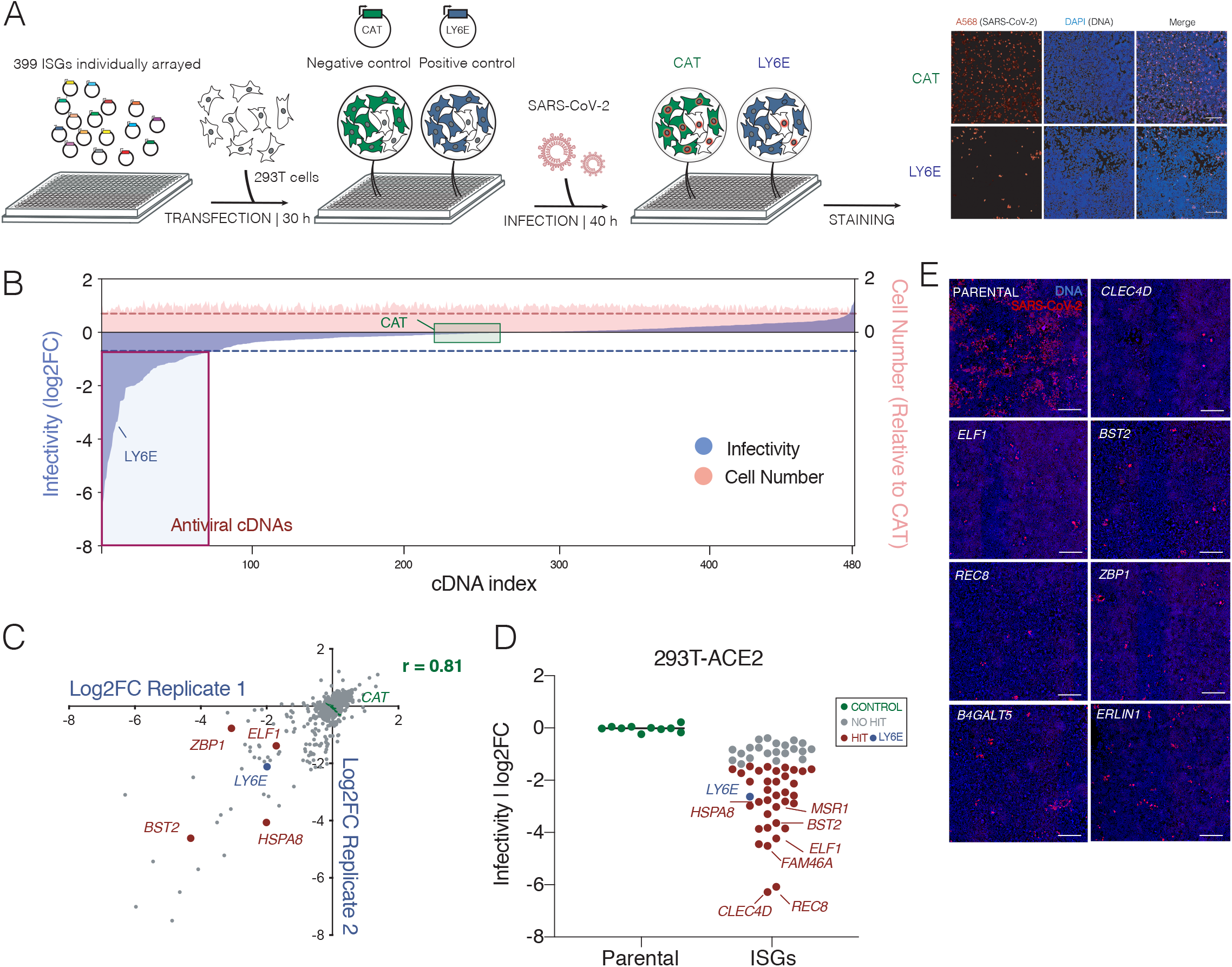
IFN-mediated restriction of SARS-CoV-2 relies on a limited subset of ISGs. (A) Schematic representation of the gain-of-function screen to identify ISGs that inhibit SARS-CoV-2 replication. (B) Ranked log2FC SARS-CoV-2 infectivity values (blue shading) and normalized cell number (pink shading), after individual overexpression of 399 human ISGs and controls. Dashed lines illustrate cut offs for antiviral ISG hit calling strategy: dotted blue line, infectivity = 4*Stdev log2FC *CAT*; dotted pink line, cell viability = 70% of *CAT*. Controls are shown (*CAT*, negative; *LY6E*, positive). (C) Correlation plots of log2FC infectivity values for ISG overexpression screens using 293T cells infected with SARS-CoV-2. r = Pearson correlation coefficient between screens. (D) 293T-ACE2 stably expressing each of the identified ISGs were infected with SARS-CoV-2 (MOI 0.25). At 40 h post-infection, cells were fixed, stained with DAPI and immunolabelled with anti-SARS-CoV-2 nucleoprotein (N) antibody. Log2FC infectivity was calculated as the percentage of N^+^/DAPI^+^ cells relative to parental control wells. Data represent mean ± SD of three independent experiments (n=3). Statistical significance was calculated using one-way ANOVA with Sidak’s multiple comparison post-hoc test. Representative images are shown in (E). Scale bar = 10 μm.

To further validate the antiviral activity of the ISGs identified in this high-throughput screen, we generated stable cell lines expressing each of these 65 ISGs and assessed their ability to inhibit SARS-CoV-2 replication. Upon transduction of 293T-ACE2 cells with lentiviruses carrying these 65 factors, 7 cell lines did not survive antibiotic selection, so stable lines could only be generated for the remaining 58 ISGs. Next, the ability of these ISG-expressing cells to support replication of SARS-CoV-2 was evaluated. Of these, 37 lines showed statistically significant reductions of SARS-CoV-2 replication compared to parental cells (log2FC at least four standard deviations lower than parental cells and p value ≤ 0.05) (**Fig 1D, E**).

Importantly, this screening approach captured both upstream regulators as well as downstream effectors of the IFN response, including the signaling adaptor *MYD88*, signal transducers *STAT1* and *STAT2*, transcription factors *ELF1, REC8*, and *ETV6*, and several IFN effectors including *BST2, IFITM2*, and *IFITM3*, which likely harbor direct antiviral activities. The full list of identified ISGs and their activities are shown in **Table S2 and S3**.

### Network model of SARS-CoV-2 antiviral effectors

ISGs are a heterogenous group of genes with encoded functions ranging from inflammatory pathway signaling to intracellular trafficking, energy metabolism, and nuclear transport (Schoggins, 2019). To better understand the biochemical and functional context by which these 65 ISGs exert antiviral activities, we conducted a supervised network propagation leveraging high confidence protein-protein interactions and hierarchical relationships (**Fig 2**, *see Methods*). Using this analysis, we identified densely interconnected protein clusters that are significantly associated with cellular biological processes (Raudvere et al., 2019). As expected, we found strong association to pathways that stimulate IFN signaling, including cytosolic pattern recognition receptors and regulators of STAT phosphorylation, as well as pathways linked to the type I IFN response, the cellular response to viral infection, and cytokine signaling (**Fig 2A**, blue boxes). We also observed an enrichment of RNA helicases, and regulators of cell death. Within this group were *DDX60*, which exhibits antiviral activity against hepatitis C virus (HCV) and VSV (Schoggins et al., 2011), *ZBP1*, which was recently identified as a sensor of influenza A virus Z-RNA motifs, and *MKLK*, a ZBP1 binding partner and downstream activator of necroptosis in response of viral infection (**Fig 2B**) (Zhang et al., 2020). Additional enriched clusters included regulation of transport at the Golgi network or the ER (**Fig 2C, D**), nucleotide metabolism, and regulators of sphingolipid metabolism, including the ISGs *B4GALT5* and *ST3GAL4* (**Fig 2E**). Additional ER/Golgi resident factors identified as potent restrictors of SARS-CoV-2 replication included the apolipropotein *APOL2* and *RSAD2*/*Viperin*, which are involved in lipid synthesis and mobilization. This suggests that regulation of the membrane composition at sites relevant for viral replication or trafficking is likely a critical host strategy for the control of SARS-CoV-2 replication. Overall, this network analysis underscores the diversity of activities that underlie the cellular antiviral response to SARS-CoV-2 replication.

**Figure 2.**
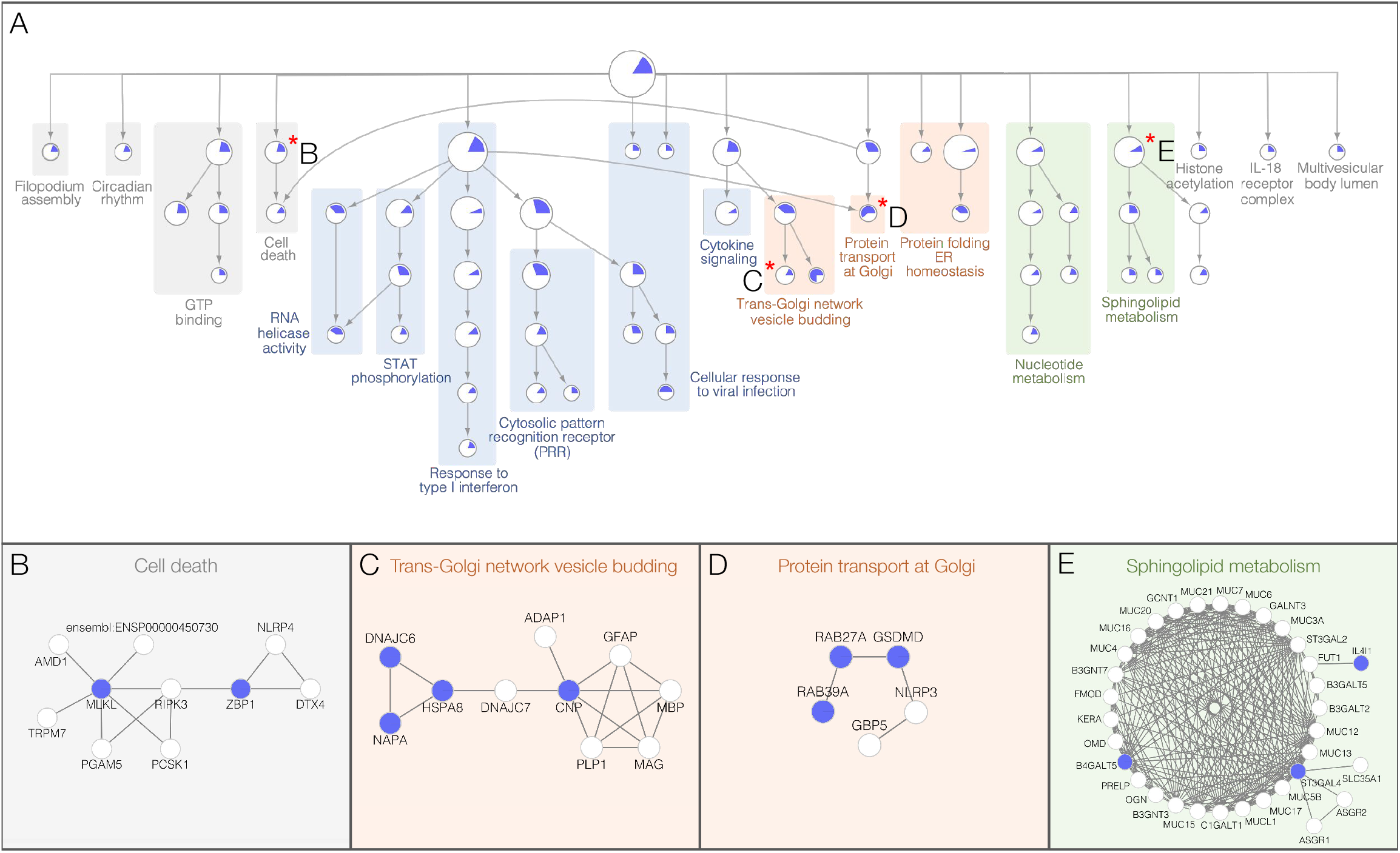
Network model of SARS-CoV-2 antiviral effectors. (A) The network containing the 65 identified antiviral ISGs was expanded to include a total of 343 high confidence protein interactors (Score> 0.7 STRING) and subjected to supervised community detection (Carlin et al., 2017; Shannon, 2003). The resultant hierarchy is shown. Here, each node represents a community of densely interconnected proteins, and each edge (arrow) denotes containment of one community (edge target) by another (edge source). Enriched biological processes are indicated. The percentage of each community that corresponds to the 65 antiviral ISGs is shown in dark blue. (B-E) Zoom-in insets from selected protein communities are indicated with an asterisk * in the hierarchy. Nodes indicate proteins, and edges indicate interactions from STRING. Blue nodes indicate ISGs that restricted SARS-CoV-2 replication.

### Restriction of SARS-CoV-2 entry

To understand how these antiviral effectors impact viral replication, a selected subset of ISGs were tested for their ability to inhibit specific stages of the SARS-CoV-2 infectious cycle. Firstly, we adopted a pseudotyped VSV expressing SARS-CoV-2 S protein (VSV-S-luciferase) to measure *viral entry* (**Fig 3A**, diagram). Then we assessed *viral RNA replication* by measuring viral RNA at 8 h post-infection (**Fig 3B**). Lastly, we infected naïve cells with viral supernatants that were collected at 18 h post infection to assess *late stage activity*, encompassing viral translation and egress (**Fig 3C**). These experimental data were integrated with available bioinformatic resources that provided information on subcellular localization and known function to establish a predictive map of the impact of these ISGs on the SARS-CoV-2 infectious cycle (**Fig 4**).

**Figure 3.**
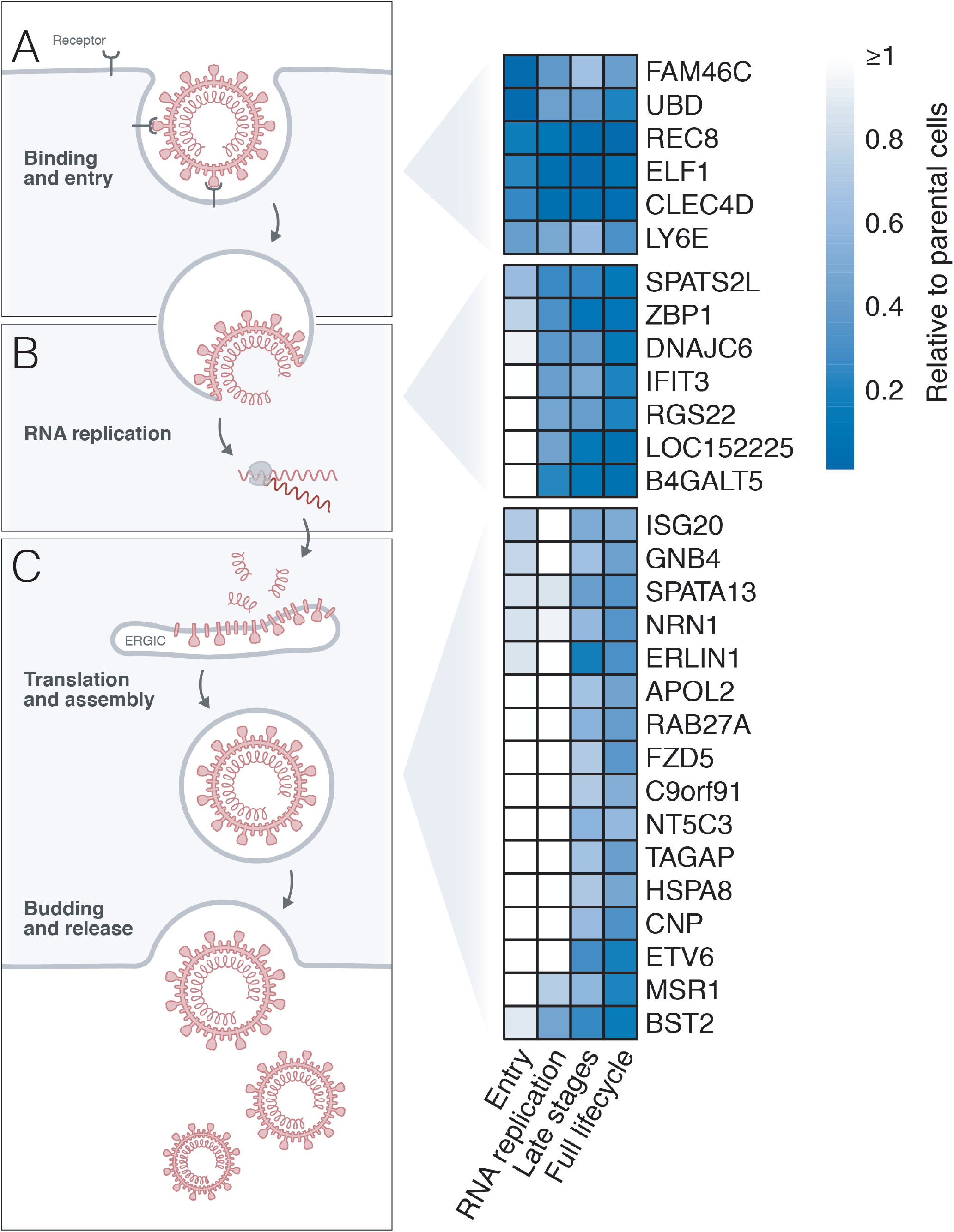
SARS-CoV-2 antiviral effectors inhibit discrete viral replication steps. 293T-ACE2 cells stably expressing each of the indicated ISGs were subjected to (A) infection with SARS-CoV-2 pseudotyped VSV luciferase virus (VSV-S-luc) for 16 h prior to measurement of luciferase signal. In parallel, cells were subjected to synchronized infection with SARS-CoV-2 (MOI = 4) for 6 h prior to measurement of viral RNA (B), or supernatants at 18 h post-infection were used to infect naïve Vero E6 cells. Infectivity was then determined at 18 h post-infection using immunostaining for viral N protein (C). In parallel to these experiments, the impact of these ISGs in 293T-ACE2 cells on SARS-CoV-2 replication at 24 h post-infection was evaluated (full lifecycle). Results are summarized in the heat map and show the mean (n=2) of relative activities compared to parental cells.

**Figure 4.**
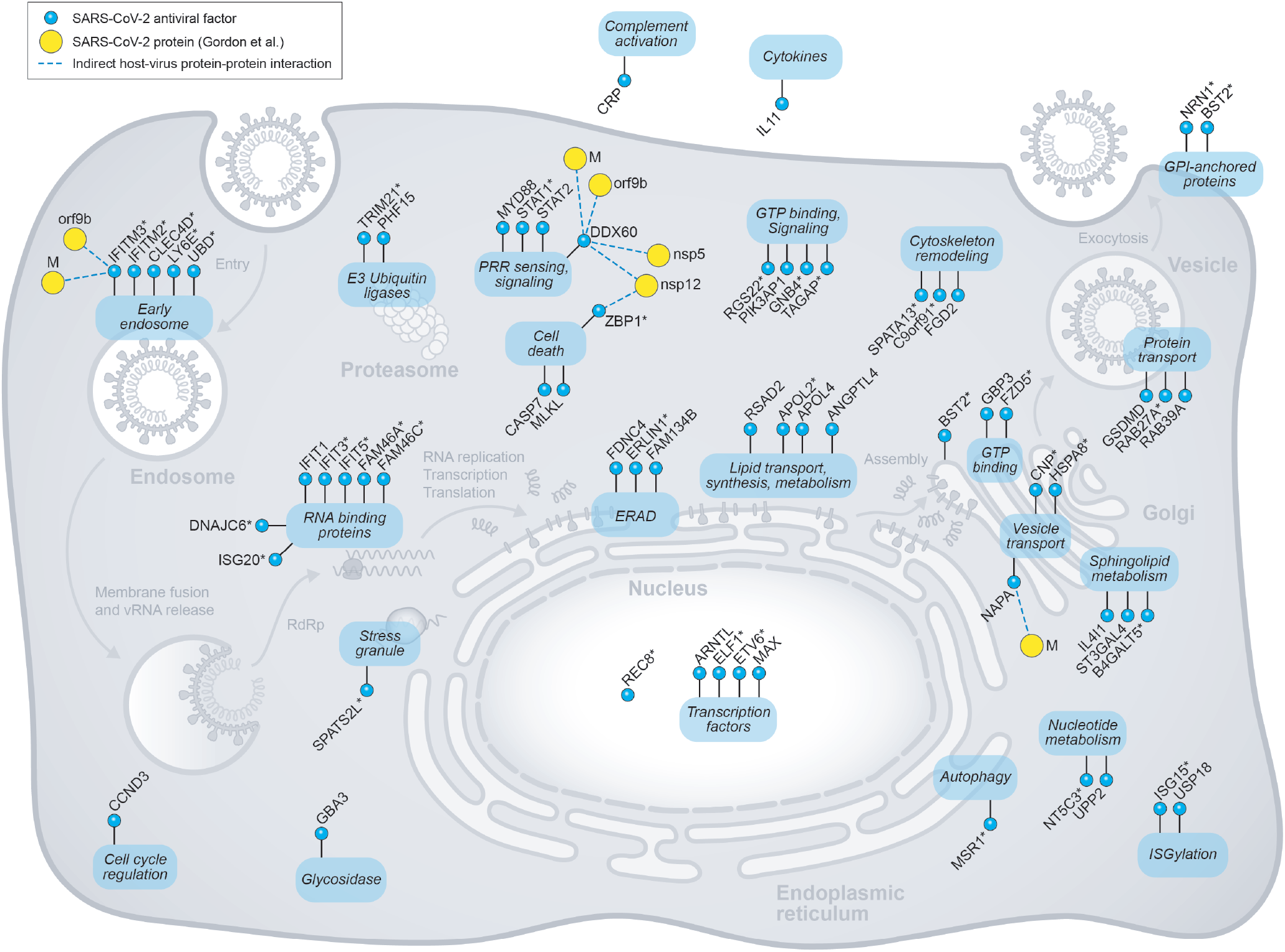
Integrated model of SARS-CoV-2 cellular restriction mechanisms. ISGs that inhibited SARS-CoV-2 replication were placed at specific positions along the viral infectious cycle based on experimental data generated in Figure 3 in conjunction with Gene Ontology, KEGG, Reactome databases and the literature. Human ISGs are represented in blue, and SARS-CoV-2 proteins in yellow. Asterisks * indicate ISGs that were validated using lentiviral transduction. Dashed lines (edges) represent indirect interactions between these ISGs and the indicated viral proteins based on constitutively expressed interactors of ISGs (Hubel et al., 2019) and reported SARS-CoV-2 interactors (Gordon et al., 2020).

Entry of SARS-CoV-2 into the host cell is facilitated by viral S protein binding to the ACE2 cellular receptor promoting endocytosis. Upon entry, SARS-CoV-2 viral particles escape the endosome to initiate viral replication (Hoffmann et al., 2020). Six ISGs reduced entry of the pseudotyped VSV-S by more than 50%, including *LY6E, CLEC4D, UBD, ELF1, FAM46C* and *REC8* (**Fig 3A, 4**). *LY6E* was previously demonstrated to restrict SARS-CoV-2 entry by inhibiting viral S protein fusion at the membrane (Pfaender et al., 2020). *CLEC4D* is an integral membrane protein that acts as an endocytic receptor and has been linked to inhibition of bacteria uptake (Wilson et al., 2015). Another ISG affecting viral entry was *UBD*/*FAT10*, which is recruited to the incoming Salmonella-containing vacuole (SCV) together with the autophagy cargo receptor p62 (Spinnenhirn et al., 2014), and these serve as signals for lysosomal targeting and pathogen clearance. Since these ISGs have been reported to impact endo-lysosomal function, it is possible that they interfere with SARS-CoV-2 entry by impeding low pH-dependent entry or endosomal escape. Finally, the transcription factor *ELF1* was also found to affect viral entry. *ELF1* governs a complex transcriptional program of over 300 genes that are largely distinct from those induced by IFN, suggesting that a secondary antiviral transcriptional cascade acts to inhibit SARS-CoV-2 entry, and potentially other stages of the viral life cycle (Seifert et al., 2019).

### Cellular inhibition of SARS-CoV-2 RNA replication

Following SARS-CoV-2 release into the cytosol, expression of the replicase gene from the viral genomic RNA generates non-structural proteins (nsp). These nsps coordinate the assembly of the replicase-transcriptase complex (RTC) at the ER, which enables viral RNA replication and protein synthesis (Fehr and Perlman, 2015). Seven ISGs were found to strongly inhibit SARS-CoV-2 RNA replication (>50% inhibition) (**Fig 3B)**, including *IFIT3, SPATS2L, DNAJC6, RGSS2, LOC152225*, as well as *ZBP1* and *B4GALT5*, which were found to be core components of the cell death and sphingolipid metabolism networks shown in **Fig 2B, E**.

The IFIT-family includes five members (*IFIT1, IFIT1B, IFIT2, IFIT3*, and *IFIT5*), which prevent active viral RNA replication by detection and sequestering of single-stranded 5′-ppp or 2′O-unmethylated RNA (Metz et al., 2013). In this study, we identified three members of this family, *IFIT1, IFIT3*, and *IFIT5*, to inhibit SARS-CoV-2 replication, suggesting this family plays an important role in the restriction of SARS-CoV-2. RNA replication was also reduced by the RNA binding protein *SPAT2SL*. Following stress stimuli, SPAT2SL is recruited to cytoplasmic stress granules, where viral RNA can be sequestered to reduce viral genome synthesis (Miller, 2011; Zhu et al., 2008). Finally, the ISG *DNAJC6*, a member of the heat shock protein 40 (HSP40) family, was also determined to impact the SARS-CoV-2 replicative stage (**Fig 3B)**. *HSP40* family members are known to play critical roles in protein transport, folding, and structural disassembly, and can bind the 3’ untranslated region of the mouse hepatitis virus (MHV) coronavirus (Nanda et al., 2003; Rosenzweig et al., 2019). Overall, these data suggest that molecular recognition and targeting of viral RNA is a critical host defense strategy used to interfere with SARS-CoV-2 genome synthesis.

### ER- and Golgi resident ISGs inhibit late stage SARS-CoV-2 replication

Transcription and translation of SARS-CoV-2 subgenomic mRNAs at the ER membrane generate accessory, as well as the structural proteins S, envelope (E), membrane (M), and nucleocapsid (N). S, E, and M are then inserted into the ER and transit through the secretory pathway to commence viral assembly in the ER–Golgi intermediate compartment (ERGIC). Specifically, M, S and E associate with viral genomes encapsidated by the N protein to form virions that bud from the ERGIC. Virions traffic in vesicles through the *trans*-Golgi network and are subsequently released by exocytosis. Notably, we found that a majority of ISGs in our assay (16/35, 55%) restricted late stages of viral replication (**Fig 3C**). Based on their reported function, late stage ISGs were clustered with predicted impacts on translation, ERAD, and vesicle trafficking.

#### Translation

The 5’-nucleotidase family member *NT5C3*, as well as the broad spectrum antiviral *ISG20*, impacted translation or egress of SARS-CoV-2. While reported activities include regulation of nucleotide pools and RNA degradation, both factors have also been implicated in the inhibition of viral translation. Specifically, *NT5C3* was found to inhibit translation of HCV proteins, and *ISG20* was shown to interfere with translation of VSV by discriminating between self and non-self mRNAs (Metz et al., 2013; Wu et al., 2019).

#### ERAD

Accumulation of viral proteins during virion assembly at the ER-Golgi interface can trigger ERAD. Accordingly, we found the ERAD regulator *ERLIN1* to strongly attenuate late stages of SARS-CoV-2 replication. Two additional factors, *RETREG1* and *FNDC4*, also involved in this pathway with roles as a ER-phagy receptor and association with the aggresome (Wilkinson, 2019), were also found to restrict SARS-CoV-2 replication, suggesting that ERAD is a critical cellular antiviral mechanism triggered during SARS-CoV-2 infection.

#### Vesicle Trafficking

Trans-Golgi vesicle budding was found as an enriched network for the control of SARS-CoV-2 replication (**Fig 2C**). Proteins within this network include the heat shock protein *HSPA8*, and the 2’,3’-cyclic nucleotide 3’ phosphodiesterase *CNP*; both mapped to late stage viral replication (**Fig 3C**). *HSPA8* is involved in vesicle uncoating, whereas *CNP* was reported to inhibit release of human immunodeficiency virus 1 (HIV-1) (Wilson et al., 2012). Notably, these ISGs were found in a protein complex with *NAPA*, another identified restriction factor for SARS-CoV-2, and a member of the SNARE complex that functions to dock and fuse vesicles to target membranes. Finally, the GTPase *Rab27a* also impeded late stages of replication. *Rab27a* controls exocytic transport through fusion of multivesicular endosomes to the plasma membrane (Ostrowski et al., 2010), further underscoring the control of vesicular trafficking as a critical antiviral mechanism to control SARS-CoV-2 replication.

### BST2 inhibits release of SARS-CoV-2 and is antagonized by Orf7a

The bone marrow stromal antigen 2 (BST2; also known as CD317 or tetherin) was identified as a potent inhibitor of SARS-CoV-2 replication (**Fig 3C**). BST2 traffics through the ER and Golgi, and localizes at the plasma membrane and in endosomes. It has been shown to inhibit viral release of several enveloped viruses, including HIV-1, human coronavirus 229E, and SARS-CoV-1, that either bud at the plasma membrane or at the ERGIC by tethering their virions to the cell surface or intracellular membranes (Neil et al., 2008; Taylor et al., 2015; Van Damme et al., 2008; Wang et al., 2014).

BST2 restriction of SARS-CoV-2 replication was further confirmed in ACE2/TMPRSS2-expressing 293T and Huh7 cells at 24 and 48 h post-infection (**Fig 5A, B, S2A**). We next conducted loss-of-function studies in HeLa cells, since these harbor constitutive expression of BST2 (Neil et al., 2008; Van Damme et al., 2008), and found that cells depleted for BST2 released significantly more infectious viruses over time (**Fig 5C, S2B**). Overall, these data strongly support a role for BST2 in the restriction of SARS-CoV-2 replication.

**Figure 5.**
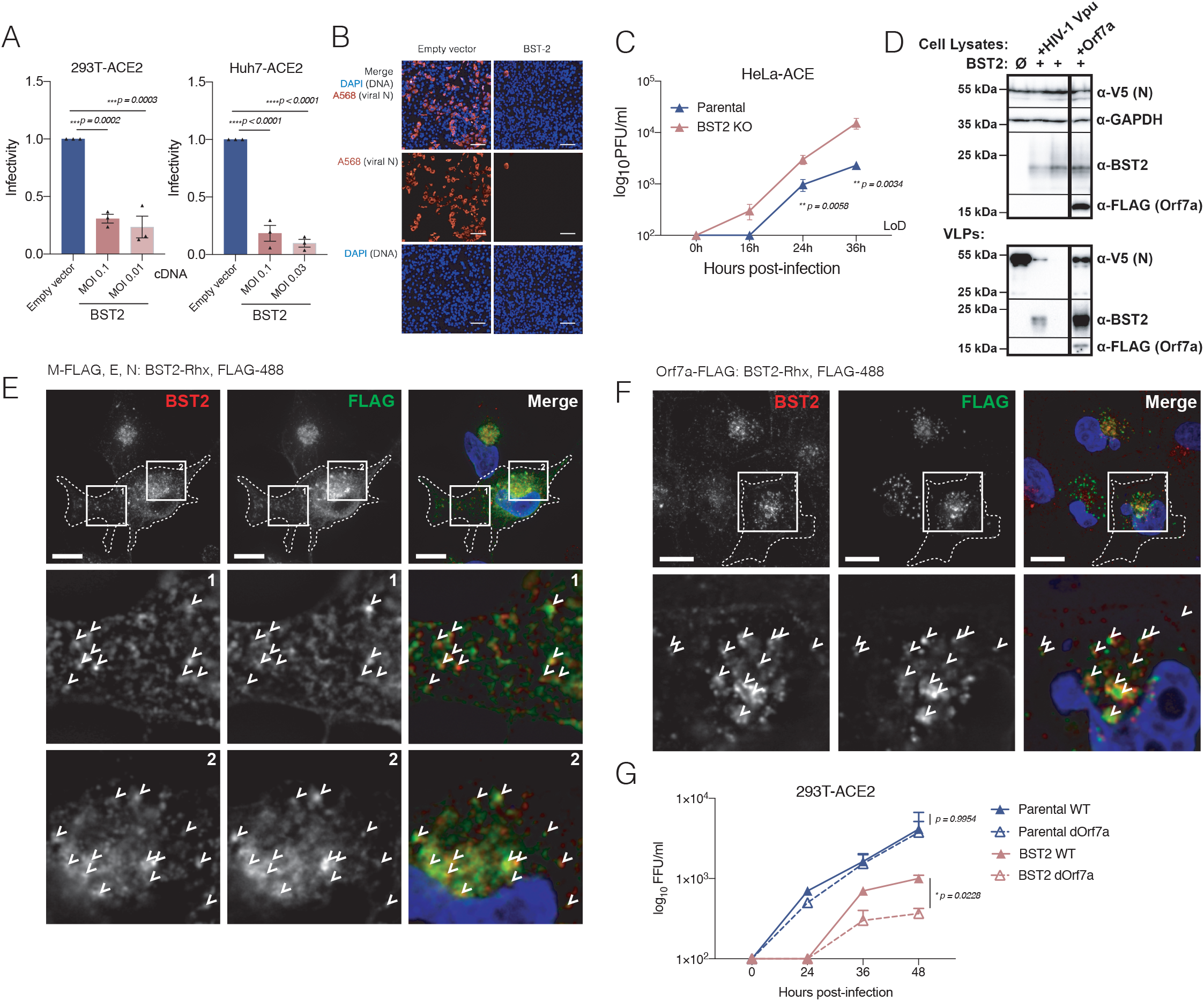
BST2 inhibits release of SARS-CoV-2 and is antagonized by Orf7a. (A, B) 293T and Huh7 cells transfected with BST2 along with ACE2 and TMPRSS2 were infected with SARS-CoV-2 at the indicated MOIs for 48 h prior to immunostaining for viral N protein. Shown is quantification of infectivity (% infected cells) relative to empty vector control (A), and representative images of Huh7 cells infected at MOI 0.03 (B). Data show mean ± SD from three independent experiments. (C) HeLa-ACE2 parental or BST2 KO cells were infected with SARS-CoV-2 (MOI = 2). At the indicated hours post-infection, supernatants were collected and analyzed by plaque assay in Vero E6 cells. LoD = limit of detection. Data show mean ± SD from one representative experiment in triplicate (n=3) of two independent experiments. (D) 293T cells were transfected with M-FLAG, E-V5, N-V5, along with the indicated constructs expressing BST2, human codon-optimized HIV-1 Vpu and/or FLAG-tagged SARS-CoV-2 Orf7a. At 24 h post-transfection, cell lysates and supernatants (VLPs) were analyzed using SDS-PAGE and immunoblotted with indicated antibodies. (E) HeLa-ACE2 cells transfected with M-FLAG, E, and N, were subjected to immunostaining for BST2 and FLAG, as indicated. Shown are deconvolved widefield microscopic images revealing colocalization of BST2 and M (arrows). Scale bar = 10 μm. (F) HeLa-ACE2 cells transfected with Orf7a-FLAG were subjected to immunostaining for BST2 and FLAG, as indicated. Shown are confocal images revealing colocalization of BST2 and Orf7a (arrows). Scale bar = 10 μm. (G) Parental 293T-ACE2 or BST2 stable cells were infected with WT or dOrf7a (MOI = 1). At indicated times post-infection, supernatants were collected and analyzed by plaque assay in VeroE6 cells. Data show mean ± SD from one representative experiment in triplicate (n=3) of two independent experiments. Statistical significance was calculated using one-way ANOVA with Dunnet’s post-hoc (A), Student’s t-test (C), or Tukey’s multicomparison test (G).

Notably, BST2 expression reduced SARS-CoV-2 RNA replication (53% reduction compared to control cells) followed by a more potent reduction of viral release (74% reduction) (**Fig 3B, C**). To further characterize the impact of BST2 on late stage replication, we evaluated viral egress in the presence or absence of BST2 using a virus-like particle (VLP) system that bypasses viral entry and viral RNA replication (Siu et al., 2008). We confirmed that this system can recapitulate virus egress, as transfection of viral M, N, and E, but not E and N alone, resulted in secreted N protein (**Fig S2C**). Using this system, we detected a strong reduction of VLP release in the presence of BST2, evidenced by loss of secreted N, corroborating that BST2 acts to inhibit egress of SARS-CoV-2 (**Fig 5D**). We next investigated if BST2 colocalizes with SARS-CoV-2 structural proteins. Notably, we detected colocalization of BST2 and structural proteins M and S at intracellular foci within the perinuclear region (caption 2, **Fig 5E, S2D**) but not at the plasma membrane (caption 1, **Fig 5E**). Together, these data indicate that BST2 spatially associates with SARS-CoV-2 structural proteins during viral assembly and trafficking.

Several viruses have developed evasion strategies to overcome restriction by BST2, including the HIV-1 accessory protein Vpu and SARS-CoV-1 Orf7a (Neil et al., 2008; Taylor et al., 2015; Van Damme et al., 2008). Notably, we found that both HIV-1 Vpu and SARS-CoV-2 Orf7a expression partially rescued BST2-mediated inhibition of SARS-CoV-2 release (**Fig 5D**), and that both Orf7a and BST2 were incorporated into the VLP particles (**Fig 5D**) (Fitzpatrick et al., 2010). We further investigated the location of BST2 and Orf7a in the cell and observed that BST2 and Orf7a colocalized in the perinuclear region (**Fig 5F**). To further investigate Orf7a antagonism of BST2, we infected parental or BST2 293T stable cells with either WT SARS-CoV-2 or a recombinant SARS-CoV-2 that was engineered to replace Orf7a with nanoluciferase (dOrf7a) (kindly provided by Ralph Baric) (Hou et al., 2020). While WT and dOrf7a viruses grew similarly in parental cells, the replication of dOrf7a virus was significantly attenuated in BST2-expressing cells at 48 h post-infection (**Fig 5G**). Overall, these data establish BST2 as a potent inhibitor of SARS-CoV-2 egress, and demonstrate that viral Orf7a protein enables immune evasion through the antagonism of BST2 restriction.

### Comparative antiviral activities of SARS-CoV-2 restriction factors

Finally, to understand if discrete cellular defense strategies are deployed to inhibit SARS-CoV-2 replication, the restriction dataset was cross-referenced with published single ISG overexpression studies that covered 20 different RNA and DNA viruses, including influenza A virus (FluAV), West Nile virus (WNV), HCV, and HIV-1 (Schoggins et al., 2011, 2014) (**Fig 6**). Interestingly, ten SARS-CoV-2 ISGs were found to reduce replication of four or more viruses (**Fig 6**). These include well described IFN signaling transducers, signaling molecules, and innate immune sensors *STAT2*, and *MYD88*, inhibitors of viral entry *IFITM2* and *IFITM3* (Brass et al., 2009), and viral nucleic acid binders *ZBP1* and *IFIT1*. Conversely, a cluster of 8 ISGs harbored selective activities for SARS-CoV-2 (**Fig 6**), including ER-Golgi resident proteins *NAPA, APOL2*, and *ERLIN1*. Notably, significant enrichment in ISGs that regulate ER homeostasis and Golgi transport suggest that these organelles are critical sites for the cellular control of SARS-CoV-2 replication. Surprisingly, many of these antiviral factors have not been reported to impact other viruses that rely on these membraned compartments for replication and assembly, including flavi-, toga-, arteri-, and bunyaviruses, suggesting that these cellular defense mechanisms target unique aspects of coronavirus translation, assembly, and egress.

**Figure 6.**
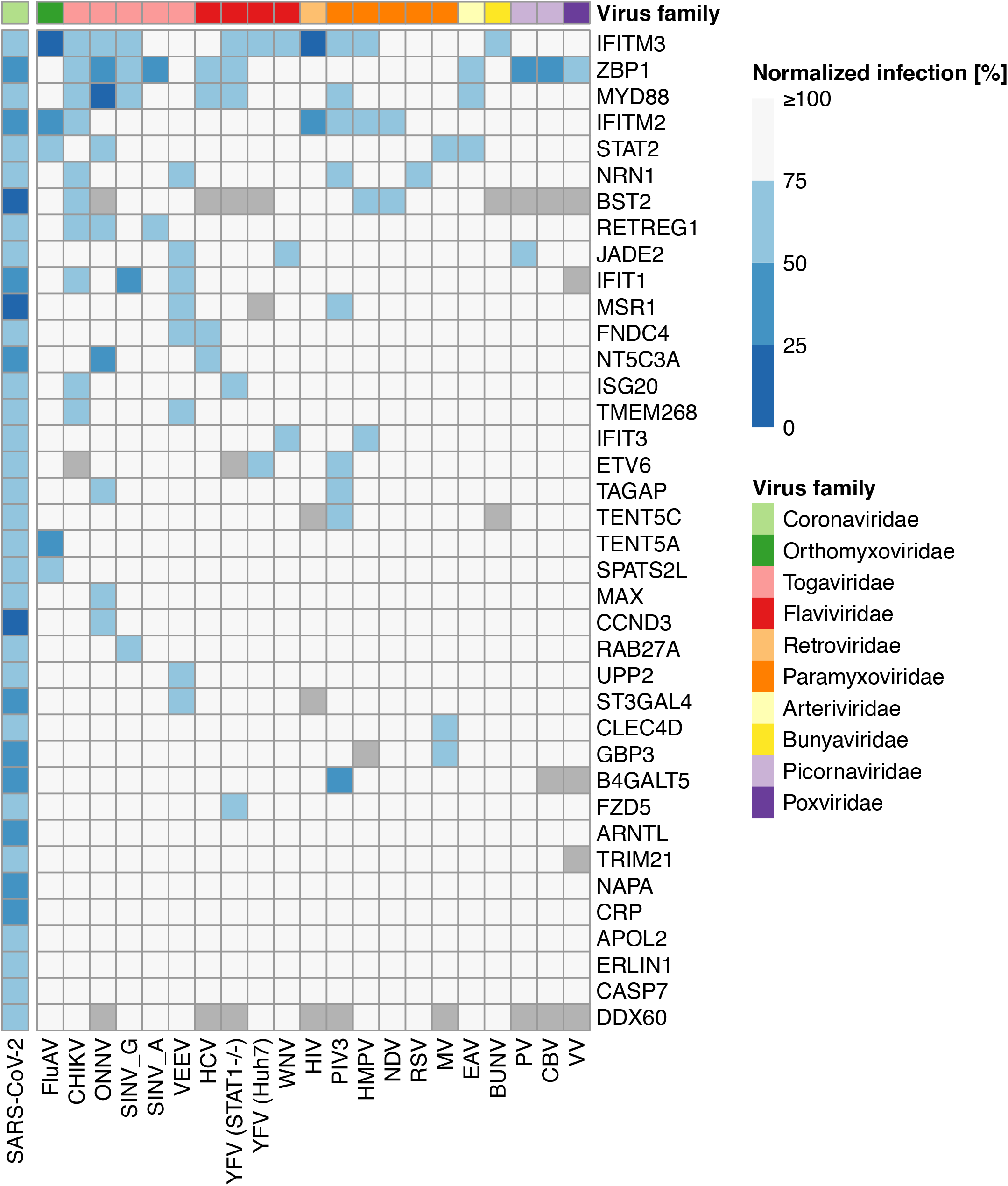
Comparative antiviral activities of SARS-CoV-2 restriction factors. Heat map showing normalized infection upon overexpression of indicated ISGs across 21 viruses. Data for SARS-CoV-2 were generated within this study. Data for the remaining 20 viruses were obtained from previously published work (Schoggins et al., 2011, 2014). Virus families are indicated. Chikungunya virus (CHIKV), O’nyong’nyong virus (ONNV), Sindbis virus (SINV), Venezuelan equine encephalitis virus (VEEV), Yellow fever virus (YFV), Human parainfluenza virus type 3 (PIV3), Human metapneumovirus (HMPV), Newcastle disease virus (NDV), Respiratory syncytial virus (RSV), Measles virus (MV), Equine viral arteritis (EVA), Bunyamwera virus (BUNV), poliovirus (PV), coxsackievirus (CBV), vaccinia virus (VV).

Taken together, this comprehensive analysis of the ISGs that act to impede SARS-CoV-2 revealed that the IFN response to SARS-CoV-2 infection relies on a limited subset of ISGs that govern a diverse set of cellular functions, including endocytosis, nucleotide biosynthesis and sphingolipid metabolism. Further dissection of these critical host-pathogen interactions, as well as potential viral evasion strategies, will enable insights into the molecular determinants of innate immune control of SARS-CoV-2 replication and clinical disease outcomes.

## DATA AVAILABILITY

The A549 and HTBE RNA-seq data used in this study have been deposited in the Gene Expression Omnibus (GEO) database repository under the accession number GSE156295 (token whyhqquupvadjgn).

## ACKNOWLEDGMENTS

We would like to thank Ralph Baric for providing the dOrf7a SARS-CoV-2, Thomas Rogers for providing the HeLa-ACE2 cells, Kwok-Yung Yuen for providing the rabbit-anti-SARS-CoV-2 N antibody, and Stefan Pohlmann for providing the mammalian expression vector encoding SARS-CoV-2 S, pCG1-CoV2-S-HA. We also would like to thank Marisol Chacon for administrative support, Sylvie Blondelle and Larry Adelman for biosafety support, and Rowland Eaden for shipping assistance. We would also like to thank the Viral Vector Core Facility at the SBP for the cDNA normalization and the lentivirus production. This work was supported by the following grants to the Sanford Burnham Prebys Medical Discovery Institute and the Icahn School of medicine at Mount Sinai: DoD: W81XWH-20-1-0270; DHIPC: U19 AI118610; Fluomics/NOSI: U19 AI135972. This work was also supported by generous philanthropic donations from Dinah Ruch and Susan & James Blair, from the JPB Foundation, the Open Philanthropy Project (research grant 2020-215611 (5384)) and anonymous donors. Additional support has been provided by DARPA grant HR0011-19-2-0020 and by CRIP (Center for research on Influenza Pathogenesis), a NIAID-funded Center of Excellence for Influenza Research and Surveillance (CEIRS, contract # HHSN272201400008C). This work was additionally supported by the following grants to Northwestern University Feinberg School of Medicine: a CTSA supplement to NCATS: UL1 TR002389; a CTSA supplement to NUCATS with the generous support of the Dixon family: UL1 TR001422; and a Cancer Center supplement: P30 CA060553, and the following grant to JG at UC San Diego: NIH grant R37AI081668. This work was also supported by a generous grant from the James B. Pendleton Charitable Trust. The funding sources had no role in the study design, data collection, analysis, interpretation, or writing of the report. The content of this study is solely the responsibility of the authors and does not necessarily represent the official views of the National Institutes of Health.

## AUTHOR CONTRIBUTIONS

L.M.-S., M.K.L., C.A.S., J.G. and S.K.C., conceived and designed the experiments. L.M.-S., M.K.L., X.Y., C.A.S., A.P.G., P.D.J, C.N., Y.P., and A.L.O., conducted and/or analyzed the experiments. L.P. and A.M.O. conducted data analysis and representation. A.R.F., M.U., M.C. and C.B. performed and/or analyzed the RNAseq experiments. D.P., C.C., S.L., B.R, and T.I. generated the network model. L.M. and J.F.H. generated essential reagents. L.M.-S., and S.K.C. wrote the manuscript with contributions from all authors. Funding Acquisition, L.P., C.B., A.G.-S., and S.K.C.

## METHODS

### Cells and Viruses

SARS-CoV-2 USA-WA1/2020, isolated from an oropharyngeal swab from a patient with a respiratory illness who developed clinical disease (COVID-19) in January 2020 in Washington, USA, was obtained from BEI Resources (NR-52281). The recombinant dOrf7a SARS-CoV-2 was kindly provided by Ralph Baric (Hou et al., 2020). These viruses were propagated using Vero E6 cells, collected after one passage, aliquoted, and stored at −80 °C. Plaque forming unit (PFU) assays were performed to titrate the cultured virus. All experiments involving live SARS-CoV-2 followed the approved standard operating procedures of the Biosafety Level 3 facility at the Sanford Burnham Prebys Medical Discovery Institute. Vero E6 (ATCC CRL-1586), HEK293T (ATCC CRL-3216), HeLa (ATCC CRL-1586), and Huh7 (Apath LLC, Brooklyn) cells were maintained in cell growth media: Dulbecco’s modified eagle medium (DMEM, Gibco) supplemented with 10 % heat-inactivated fetal bovine serum (FBS, Gibco), 50 U/mL penicillin - 50 µg/mL streptomycin (Fisher Scientific), 1 mM sodium pyruvate (Gibco), 10 mM 4-(2-hydroxyethyl)-1-piperazineethanesulfonic acid (HEPES, Gibco), and 1X MEM non-essential amino acids solution (Gibco). BHK-21/WI-2 cells (Kerafast, MA) were maintained in DMEM (Gibco) supplemented with 10 % heat-inactivated FBS (Gibco) and 50 U/mL penicillin - 50 µg/mL streptomycin. Human tracheobronchial epithelial (HTBE) cells (ATCC PCS-300-010) were cultured in commercially available airway epithelial cell basal medium following manufacturer’s protocol (ATCC). HTBE cells were derived from one donor and all tissues used for isolation of these cells were obtained under informed consent and conform to HIPAA standards to protect the privacy of the donors’ personal health information. HEK293T and HeLa cells stably expressing ACE2 (293T-ACE2/HeLa-ACE2) were generated by transducing HEK293T or HeLa cells with human ACE2-expressing lentiviruses, followed by selection of resistant cells with puromycin (InvivoGen) at 2 μg/ml for 14 days. The resistant cells were then maintained in cell growth media supplemented with 1 μg/ml puromycin. ACE2 expression was confirmed by western blot analysis. All cells were tested and were confirmed to be free of mycoplasma contamination.

### Antibodies

The antibodies used in this study include: *Immunofluorescence:* rabbit-anti-SARS-CoV-2 N antibody (gift from Kwok-Yung Yuen, University of Hong Kong), mouse anti-HM1.24 (BST2) (a gift from Chugai Pharmaceutical Co., Kanagawa, Japan), rat anti-FLAG-AlexaFluor-488 (Biolegend, #637317), mouse anti-HA-AlexaFluor-594 (Biolegend, #901511), donkey anti-mouse-AlexaFluor-488 (Jackson ImmunoResearch, #715-545-150), donkey anti-mouse-Rhodamine-Red-X (Jackson ImmunoResearch, #715-295-150). *Western blotting:* rabbit polyclonal anti-BST2 (NIH AIDS Reagent Program, Division of AIDS, NIAID, NIH: Anti-BST-2 Polyclonal (cat# 11721) from Drs. Klaus Strebel and Amy Andrew), mouse monoclonal anti-V5 tag (Invitrogen, #R960-25), mouse monoclonal anti-GAPDH (GeneTex, #GTX627408), mouse monoclonal anti-FLAG M2 (Sigma, #F1804), rabbit monoclonal anti-β-actin antibody (Cell Signaling, #4970) and rabbit monoclonal anti-CoxIV antibody (Cell Signaling #4850).

### Plasmids

*Lentiviral constructs:* pLX304 constructs for each of the ISGs, and GFP and CAT as negative controls were obtained from the ORFeome library. psPAX2 (Addgene, #12260), and pMD2.G (Addgene, #12259). *SARS-CoV-2 contructs:* dsDNA gene fragments (gBlocks) encoding human-codon-optimized SARS-CoV-2 proteins M, M-FLAG, E, E-V5, N, N-V5, and Orf7a N- or C-terminally tagged with 3xFLAG tag, corresponding to the SARS-CoV-2 Wuhan-Hu-1 isolate (genbank MN908947.3), were synthesized by Integrated DNA Technologies (IDT). The gene fragments were inserted into the pcDNA3.1(-) backbone between NotI and EcoRI restriction sites using an In-fusion seamless cloning strategy (Takara Bio). The mammalian expression vector encoding COV2 S, pCG1-COV2-S-HA, was obtained from Prof. Stefan Pohlmann (Infection Biology Unit, German Primate Center - Leibniz Institute for Primate Research, University Göttingen (Hoffmann et al., 2020)).

### RNA-seq experiments

HTBE and A549 cells were seeded overnight and then treated with 100 IU/ml universal interferon beta (IFN, R&D Systems), or left untreated. At 8 h post-treatment, cell were lysed in Trizol (Thermo Fisher) and RNA was extracted using RNeasy Mini Kit (Qiagen). Strand-specific ribosomal RNA-depleted sequencing libraries were produced according to standard Illumina protocols, and sequencing was carried out on an Illumina HiSeq 2500. The human hg38 reference genome and RefSeq gene annotation were used for spliced read alignment and gene assignment. Experiments were conducted in duplicate and 412 genes were defined as ISGs based on log2FC > 1.5 and p value < 0.05.

### Overexpression cDNA screen

A targeted overexpression cDNA screen was carried out in human epithelial cells to identify ISGs that restrict the replication of SARS-CoV-2. 399 ISGs were selected for this gain-of-function screen based on experimental, published data, and availability as full-length, sequence-validated cDNA clones. These cDNAs were hand-picked from the ORFeome collection, which contains ∼17,000 full-length, sequenced, V5-epitope tagged human ORFs in the lentiviral expression vector pLX304. Each of these 399 cDNAs were individually arrayed in 384-well plates at a concentration of 40ng/well along with human ACE2 and TMPRSS2 (10 ng), and 0.25 μl of the transfection reagent Fugene 6 (Promega). After 20 min incubation at room temperature, 3,000 293T cells diluted in cell growth media (see *cells and viruses* section) were seeded per well and incubated at 37°C, 5% CO2. At 30 h post-transfection, cells were mock-treated or infected with SARS-CoV-2 (USA-WA1/2020) at a MOI 0.0625 for 40 h at 37°C, 5% CO2. Cells were then fixed with 5% PFA (Boston BioProducts) for 4 hours at room temperature and then washed twice with 1xPBS. Cells were permeabilized with 0.5% Triton X-100 for 20 min, followed by two washes with 1xPBS and blocking with 3% BSA (Sigma) for 1 h at room temperature. Anti-SARS-CoV-2 N rabbit serum was added for 1 h at room temperature, followed by three washes with 1xPBS and a 1-h incubation with Alexa Fluor 568-conjugated anti-rabbit secondary antibody (Thermo Fisher Scientific) diluted in 3% BSA. Following three washes with PBS, cells were stained with DAPI (4,6-diamidine-2-phenylindole, KPL), and plates were sealed and stored at 4°C until imaging.

### High-content imaging and data analysis

Viral replication was assessed using high-throughput microscopy. The assay plates were imaged using the IC200 imaging system (Vala Sciences) located at the Conrad Prebys Center for Chemical Genomics (CPCCG). The analysis software Columbus v2.5 (Perkin Elmer) was used to calculate infectivity (number Alexa 568+ objects/number DAPI+ objects). Screens were run in duplicate and the infectivity values for each well were normalized to the median of the negative control CAT, and used to calculate the log2FC. The hit calling strategy was based on log2FC. Factors with a corresponding log2FC < 4*Stdev CAT, and cell viability > 70 % CAT were considered restriction factors.

### Generation lentivirus and 293T-ACE2-ISG/GFP cells

Lentiviruses were generated for each of the 65 ISGs that were found to restrict SARS-CoV-2 replication. Briefly, 293T cells at passage 10 were cultured in monolayer on matrigel-coated plates. After reaching 90% of density, three plasmids, including pLX304-ISG/GFP, psPAX2 (Addgene), and pMD2.G (Addgene), were co-transfected into cells at a ratio of 3:2:1 using PEI (VWR). After 16 h incubation, transfection media were replaced with fresh DMEM supplemented with 10% FBS. Viral supernatants were collected at 48 h post-transfection with an estimated transduction unit of 2x 10^4^ lentiviral particles. Lentiviruses were used to transduce 293T-ACE2 cells (MOI = 3) pre-treated with 10 μg/ml Polybrene (Life Technologies), followed by selection of resistant cells with Blasticidin (InvivoGen) at 10 μg/ml for 14 days. 293T-ACE2-ISG/GFP resistant cells were maintained in cell growth media supplemented with 2 μg/ml Blasticidin.

### Network analyses

To understand the biochemical and functional context by which the identified antiviral ISGs function, we explored a network-based approach that could integrate these ISGs (“seed” proteins) with existing knowledge. Towards this aim, we used a pipeline that employs a combination of scripts and Cytoscape applications. First, to explore the highest confidence interactions of “seed” proteins, we selected the STRING - Human Protein Links - High Confidence (Score >= 0.7) protein-protein interaction network available on NDEx as the “background” network (link provided below). We then performed network propagation to select a neighborhood of 343 proteins ranked highest by the algorithm with respect to these seeds (Carlin et al., 2017). This “neighborhood” network (including all edges among the 343 proteins) was extracted from the background network. We then identified densely interconnected regions, i.e. “communities” within the neighborhood network, using the community detection algorithm HiDeF via the Community Detection APplication and Service (CDAPS) (manuscript in press, app available at http://apps.cytoscape.org/apps/cycommunitydetection). The result of HiDeF from CDAPS was a “hierarchy” network where each node represented a community of proteins, and edges denoted containment of one community (the “child”) by another (the “parent”). Finally, the hierarchy network was styled, communities were labeled by functional enrichment using gProfiler (via CDAPS) and a layout was applied. The STRING - Human Protein Links - High Confidence (Score > = 0.7) network is available in the Network Data Exchange (NDEx) at http://ndexbio.org/#/network/275bd84e-3d18-11e8-a935-0ac135e8bacf.

### Generation pseudotyped SARS-CoV-2 virus

VSV pseudotyped with spike (S) protein of SARS-CoV-2 was generated according to a published protocol (Whitt, 2010). Briefly, BHK-21/WI-2 cells (Kerafast, MA) transfected with SARS-CoV-2 S protein were inoculated with VSV-G pseudotyped ΔG-luciferase VSV (Kerafast, MA). After a 2 hour incubation at 37 °C, the inoculum was removed and cells were treated with DMEM supplemented with 5 % FBS, 50 U/mL penicillin, and 50 µg/mL streptomycin. Pseudotyped particles were collected 24 h post-inoculation, then centrifuged at 1,320×g to remove cell debris and stored at −80 °C until use.

### Mapping into SARS-CoV-2 infectious cycle studies

Mapping studies were conducted in parallel using 293T-ACE2-ISG/GFP cells. Briefly, multiple 96-well plates were seeded with 50,000 293T-ACE2-ISG/GFP cells/well and incubated overnight at 37°C, 5% CO2. To determine the effect of the identified ISGs on *viral entry*, 293T-ACE2-ISG/GFP cells were infected with VSV-S-luciferase and incubated for 16 h. The activity of firefly luciferase was then measured using the bright-Glo™ luciferase assay (Promega) for quantitative determination. To measure *RNA replication* and *late stages*, cells were infected with SARS-CoV-2 (USA-WA1/2020) at a MOI 4 for 1 h on ice. Viral inoculum was removed and cells were washed twice with 1xPBS and supplemented with cell growth media (see *cells and viruses* section). At 6 h post-infection, SARS-CoV-2 *RNA replication* was measured. Briefly, intracellular viral RNA was purified from infected cells using the TurboCapture mRNA Kit (Qiagen) in accordance with the manufacturer’s instructions. The purified RNA was subjected to first-strand cDNA synthesis using the high-capacity cDNA reverse transcription kit (Applied Biosystems, Inc). Real-time quantitative PCR (RT-qPCR) analysis was then performed using TaqPath one-step RT-qPCR Master Mix (Applied Biosystems, Inc) and, ActinB CTRL Mix (Applied Biosystems, Inc) for housekeeping genes, and the following primers and probe for qPCR measurements of viral genes: N-Fwd: 5’-TTACAAACATTGGCCGCAAA-3’; N-Rev: 5’-GCGCGACATTCCGAAGAA-3’; N-Probe: 5’-FAM-ACAATTTGCCCCCAGCGCTTCAG-BHQ-3’. To evaluate *late stages*, supernatants collected at 18 h post-infection were used to infect naïve Vero E6 cells. At 18 h post-infection cells were then fixed with 5% PFA (Boston BioProducts) for 4 hours at room temperature and then subjected to immunostaining and imaging for SARS-CoV-2 N protein and DAPI (described in *overexpression cDNA screen* section).

### Generation of CRISPR-Cas9 BST2 KO HeLa-ACE2 cells

Detailed protocols for RNP production have been previously published (Hultquist et al., 2019). Briefly, lyophilized guide RNA (gRNA) and tracrRNA (Dharmacon) were suspended at a concentration of 160 µM in 10 mM Tris-HCL, 150mM KCl, pH 7.4. 5µL of 160µM gRNA was mixed with 5µL of 160µM tracrRNA and incubated for 30 min at 37°C. The gRNA:tracrRNA complexes were then mixed gently with 10µL of 40µM Cas9 (UC-Berkeley Macrolab) to form CRISPR-Cas9 ribonucleoproteins (crRNPs). Five 3.5µL aliquots were frozen in Lo-Bind 96-well V-bottom plates (E&K Scientific) at -80°C until use. BST2 gene was targeted by 5 pooled gRNA derived from the Dharmacon pre-designed Edit-R library for gene knock-out. *BST2* (g1:TGCATCCAGGGAAGCCATTA, CM-011817-01; g2:TTGGGCCTTCTCTGCATCCA, CM-011817-02; g3:TTGAGGAGCTTACCACAGTG, CM-011817-03; g4: TCACTGCCCGAAGGCCGTCC, CM-011817-04; g5: CACCATCAAGGCCAACAGCG, CM-011817-05). Non-targeting negative control gRNA (Dharmacon, U-007501) was delivered in parallel. Each electroporation reaction consisted of 2.5×10^5 HeLa-ACE2 cells, 3.5 µL crRNPs, and 20 µL electroporation buffer. HeLa-ACE2 cells were grown in fully supplemented MEM (10% FBS, 1xPen/Strep, 1x non-essential amino acids) to 70% confluency, suspended and counted. crRNPs were thawed and allowed to come to room-temperature. Immediately prior to electroporation, cells were centrifuged at 400xg for 3 minutes, supernatant was removed by aspiration, and the pellet was resuspended in 20 µL of room-temperature SE electroporation buffer plus supplement (Lonza) per reaction. 20 µL of cell suspension was then gently mixed with each crRNP and aliquoted into a 96-well electroporation cuvette for nucleofection with the 4-D Nucleofector X-Unit (Lonza) using pulse code EO-120. Immediately after electroporation, 80 µL of pre-warmed media was added to each well and cells were allowed to rest for 30 minutes in a 37°C cell culture incubator. Cells were subsequently moved to 12-well flat-bottomed culture plates pre-filled with 500 µL pre-warmed media. Cells were cultured at 37°C / 5% CO2 in a dark, humidified cell culture incubator for 4 days to allow for gene knock-out and protein clearance prior to downstream applications.

### SARS-CoV-2 viral growth assays

To evaluate SARS-CoV-2 viral growth, the amount of released infectious particles was measured by plaque assay. Briefly, supernatants from SARS-CoV-2 infected cells were collected at indicated time points and stored at -80°C until used. 600,000 Vero E6 cells were seeded and incubated overnight at 37°C / 5% CO2 in 12-well plates. Confluent Vero E6 cells were then washed once with 1xPBS and infected with 100µl of virus-containing supernatants that were serially diluted 1:10. Plates were incubated 1 h at room temperature, followed by inoculum removal and addition of 1ml overlay media (2xMEM and 2.5% Avicel (FMC BioPolymer, RC-591 NF) at 1:1 ratio). 2xMEM contains 100 ml 10x MEM (Gibco), 10 ml 100x penicillin-streptomycin (Fisher Scientific), 10 ml 100x L-Glutamine, 6 ml 35% BSA, 10 ml 10 mM 4-(2-hydroxyethyl)-1-piperazineethanesulfonic acid (HEPES, Gibco), 24 mL 5% NaHCO3 (Gibco) and 340 ml water. Plates were incubated 3 days at 37°C, 5%CO2, and then fixed and stained using 0.1% Crystal Violet and 5% PFA (Boston BioProducts) overnight at 4°C.

### VLP assays

HEK-293T cells seeded in 6-well plates were transfected using Lipofectamine 2000 (Thermo-Fisher) with 625 ng each of plasmids encoding M-FLAG, E-V5, N-V5 (Fig 5D), or 500 ng of M, E, and N-V5 (Fig S2C), with or without 625 ng 3xFLAG-Orf7a or human codon-optimized HIV-1 Vpu (pVpHu from Klaus Strebel) with or without 75 ng BST2 (pcDNA3.1-BST-2 from Autumn Ruiz and Edward Stephens). After 24 hours, supernatants were collected and clarified of cell debris then pelleted through 20% sucrose at 23,500 x g for 1 hr at 4°C. Pelleted VLPs and cells were lysed in 2X Laemmli SDS-PAGE buffer, then run on 12% SDS-PAGE gels, transferred to PVDF membranes and blotted with the indicated antibodies.

### Colocalization studies

*Immunofluorescence Staining:* 2×10^4^ HeLa-ACE2 cells were seeded on 12 mm glass coverslips in 24-well plates, 24 h prior to transfection. The cells were transfected with 800 ng total plasmid DNA, using Lipofectamine 2000 (Thermo-Fisher), diluted in Optimem, according to manufacturer’s instructions. HeLa-ACE2 cells were either transfected with equal amounts (200 ng) of SARS-CoV-2 structural proteins M, E, N, S-HA, or M-FLAG, E, N, and empty plasmid (pcDNA3.1). HeLa-ACE2 cells were also transfected with 800 ng ORF7a-3xFLAG. 24 h post-transfection, cells were washed briefly in 4°C PBS before incubation with ice-cold 4% paraformaldehyde (PFA, diluted in PBS, pH 7.4). The PFA was allowed to warm to RT as the cells were fixed for 20 minutes, the PFA was removed and cells washed 3x in 1XPBS (5 min per wash). The fixed cells were quenched with 50 mM Ammonium chloride (in PBS) for 5 minutes RT, washed 3 x in PBS, and permeabilized with 0.2% Triton X-100 for 7 minutes (RT). The cells were again washed in three times in 1X PBS before incubation with 2% bovine serum albumin (BSA) in PBS for 30 minutes, prior to incubation with primary antibodies overnight at 4°C. Cells transfected with M, E, N and S-HA were stained overnight with mouse anti-HM1.24 (BST-2) antibody (diluted 1:300 in 1% BSA in PBS) at 4°C. The following day, the cells were washed 3x PBS and incubated with donkey anti-mouse-AlexaFluor-488 (1:400) for 2hr RT. The cells were washed 3x PBS (10 min per wash) and blocked with 2% BSA in PBS supplemented with 5% normal mouse serum for 1 hr RT, briefly washed in 2% BSA, and incubated with mouse anti-HA-Alexa-594 (1:200) and 4′,6-diamidino-2-phenylindole (DAPI), diluted to 1 µg/ml for 2 hr RT. Cells transfected with M-FLAG, E and N were stained overnight with mouse anti-BST-2 (diluted 1:300 in 1% BSA in PBS). The following day the cells were washed 3x PBS and incubated with donkey anti-mouse-Rhodamine-Red-X (1:400) for 2 hr RT. The cells were washed 3x PBS (10 min per wash) and blocked with 2% BSA in PBS supplemented with 5% normal mouse serum for 1 hr RT, briefly washed in 2% BSA, and incubated with rat anti-FLAG-Alexa-488 (diluted 1:200) and 1 µg/ml DAPI for 2 hr RT. Cells transfected with Orf7a-3xFLAG were stained overnight with mouse anti-HM1.24 (diluted 1:300). The following day the cells were washed 3x PBS and incubated with donkey anti-mouse-Rhodamine-Red-X (1:400) for 2 hr RT. The cells were washed 3x PBS (10 min per wash) and blocked with 2% BSA in PBS supplemented with 5% normal mouse serum for 1 hr RT, briefly washed in 2% BSA, and incubated with rat anti-FLAG-Alexa-488 (diluted 1:200) and 1 µg/ml DAPI for 2 hr RT. Following immunostaining, the cells were washed extensively in PBS, and briefly in distilled-water, before mounting in Mowiol (Polyvinyl alcohol) mounting medium (prepared in-house). *Microscopy:* Images were captured at 100x magnification (1344 ×1024 pixels) using an Olympus IX81 widefield microscope fitted with a Hamamatsu CCD camera. For each field, a Z-series of images was collected, deconvolved using the nearest-neighbor algorithm (Slidebook software V.6, Imaging Innovations, Inc) and presented as Z-stack projections. Inset images are deconvolved single z-section images. Arrow heads indicate areas of colocalization, scale bar = 10 µm. Image brightness was adjusted using Adobe Photoshop CS3.

