## Supplemental Materials for "Functional Landscape of SARS-CoV-2 Cellular Restriction"

**This PDF file includes:**

Figure legends –Figs. S1 and S2
Figs. S1 and S2
Supplementary Tables S1A, S1B, S2, and S3

**FIGURE LEGENDS**

**Figure S1 – A limited subset of ISGs account for IFN-mediated restriction of SARS-CoV-2**
(A) Volcano plots of A549 and HTBE cells stimulated with 100 IU/ml IFN for 8 h. ISGs were selected based on cut off criteria of log2FC > 1.5 and p value < 0.05. (B) Venn diagram shows overlap between ISGs identified using A549 or HTBE RNAseq datasets or previously reported (Schoggins et al., 2011). Numbers in the diagram indicate (# ISGs in the dataset/# ISGs available as validated, full-sequence length cDNA clones). (C) Overlap between 65 identified anti-SARS-CoV-2 ISGs with upregulated genes found in RNAseq analyses of SARS-CoV-2-infected *in vitro* models (Blanco-Melo et al., 2020; Emanuel et al., 2020; Overmyer et al., 2020; Sun et al., 2020).

**Figure S2 – BST2 inhibits release of SARS-CoV-2 and is antagonized by Orf7a**
(A) 293T and Huh7 cells transfected with BST2 along with ACE2 and TMPRSS2 were infected with SARS-CoV-2 at indicated MOIs for 24 h prior to immunostaining for viral N protein. Shown is quantification of infected cells relative to empty vector control, as mean ± SD from three independent experiments (n=3). Statistical significance was calculated using one-way ANOVA with Dunnet´s post-hoc. (B) Cell lysates from HeLa-ACE2 parental and BST2 KO cells were subjected to SDS-PAGE and immunoblotted using antibodies specific for BST2 and Cox-IV (loading control). (C) 293T cells were transfected with M, E, and/or N-V5. After 24 hours, cell lysates and supernatants (VLPs pelleted through 20% sucrose) were analysed by SDS-PAGE and blotted with mouse anti-V5 (for N) or mouse anti-GAPDH (loading control) antibodies. (D) HeLa-ACE2 cells transfected with M, E, N, and S-HA were subjected to immunostaining for BST2 and HA, as indicated. Shown are images revealing colocalization of BST2 and S (arrows). Scale bar = 10 µm.

Figure S1

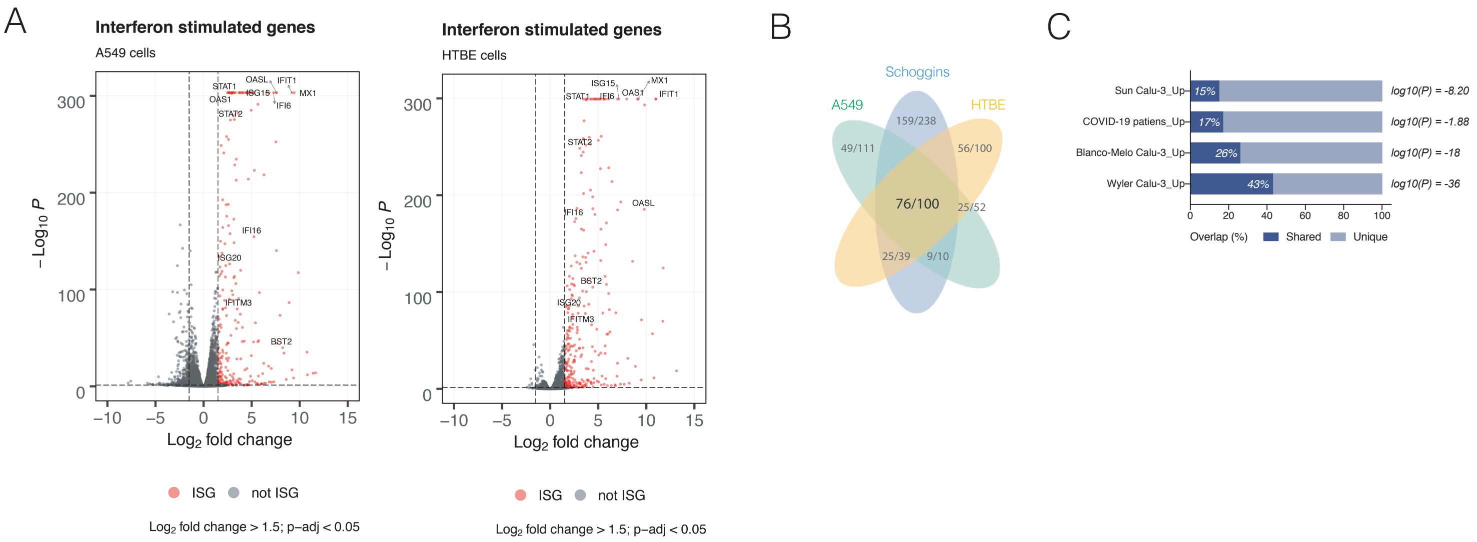

A
Interferon stimulated genes
A549 cells
Interferon stimulated genes
HTBE cells
ISG
not ISG
Log2 fold change > 1.5; p–adj < 0.05
B
Schoggins
A549
HTBE
76/100
C
Sun Calu-3_Up
COVID-19 patiens_Up
Blanco-Melo Calu-3_Up
Wyler Calu-3_Up
log10(P) = -8.20
log10(P) = -1.88
log10(P) = -18
log10(P) = -36
Overlap (%)
Shared
Unique

### Figure S2

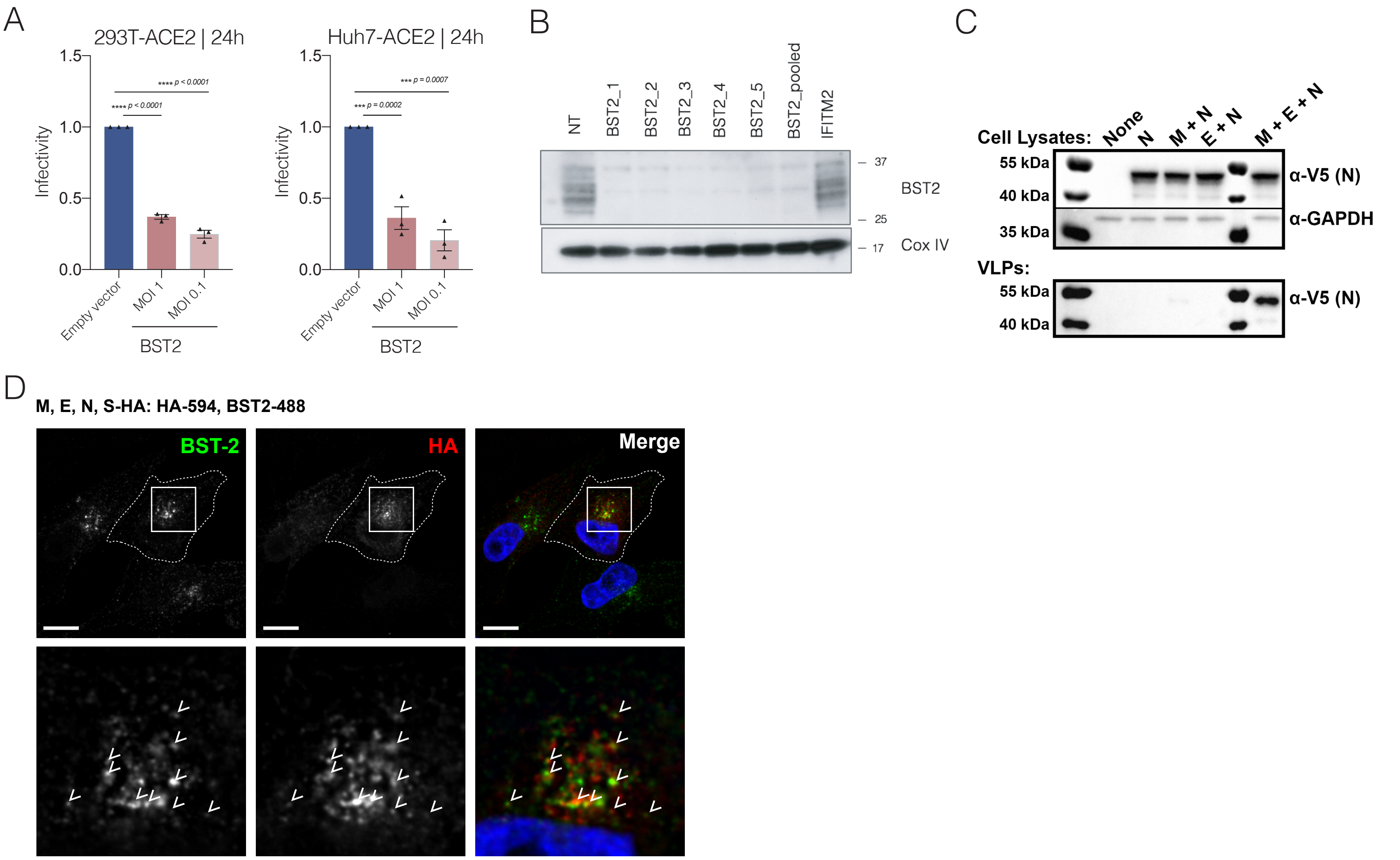

A
293T-ACE2 | 24h
Huh7-ACE2 | 24h
Infectivity
**** p < 0.0001
*** p = 0.0002
*** p = 0.0007
Empty vector
MOI 1
MOI 0.1
BST2
B
NT
BST2_1
BST2_2
BST2_3
BST2_4
BST2_5
BST2_pooled
IFITM2
BST2
Cox IV
C
Cell Lysates:
None
N
M + N
E + N
M + E + N
55 kDa
40 kDa
35 kDa
α-V5 (N)
α-GAPDH
VLPs:
D
M, E, N, S-HA: HA-594, BST2-488
BST-2
HA
Merge

**Table S1A | Identification ISGs**

**A549 and HTBE cells were treated with IFN for 8 h and subjected to RNAseq.**

Genes with log2FC > 1.5 and adjusted p value < 0.05 were considered ISGs

| Gene Symbol | Gene ID | log2FoldChange | Adjusted p-value | Cell line |
|---|---|---|---|---|
| *ACE2* | 59272 | 5.779138938 | 3.4621E-11 | HTBE |
| *ACKR4* | 51554 | 6.025002198 | 0.002694968 | A549 |
| *ACSL5* | 51703 | 2.325119652 | 0.004843387 | A549 |
| *ACSL5* | 51703 | 1.682002387 | 0.00052786 | HTBE |
| *ACY3* | 91703 | 2.386828944 | 2.94621E-05 | A549 |
| *ADAR* | 103 | 2.004721836 | 6.3097E-117 | HTBE |
| *ADGRE1* | 2015 | 2.568029692 | 1.52302E-45 | A549 |
| *AIM1* | 9212 | 2.76288684 | 1.69138E-12 | A549 |
| *AIM2* | 9447 | 7.324390028 | 1.25581E-05 | HTBE |
| *ANGPTL4* | 51129 | 2.516955983 | 2.57646E-45 | A549 |
| *APOBEC3D* | 140564 | 4.528441106 | 0.028579312 | A549 |
| *APOBEC3D* | 140564 | 1.981910987 | 0.015203147 | HTBE |
| *APOBEC3F* | 200316 | 1.611207327 | 8.46192E-07 | HTBE |
| *APOBEC3G* | 60489 | 3.058716852 | 6.34487E-52 | HTBE |
| *APOL1* | 8542 | 5.289601115 | 1.0578E-223 | A549 |
| *APOL1* | 8542 | 3.765507193 | 4.665E-101 | HTBE |
| *APOL2* | 23780 | 2.59817915 | 1.0385E-166 | A549 |
| *APOL2* | 23780 | 3.089005199 | 7.8454E-130 | HTBE |
| *APOL3* | 80833 | 6.253790508 | 1.66155E-24 | A549 |
| *APOL3* | 80833 | 5.065146935 | 1.15441E-35 | HTBE |
| *APOL4* | 80832 | 6.960351998 | 9.88685E-14 | HTBE |
| *APOL6* | 80830 | 4.002277427 | 0 | A549 |
| *APOL6* | 80830 | 3.712095145 | 1.5334E-252 | HTBE |
| *AREG* | 374 | 1.771552062 | 1.95555E-64 | A549 |
| *ARL14* | 80117 | 3.614088579 | 0.000124427 | HTBE |
| *ARRDC3* | 57561 | 1.665288035 | 6.09919E-45 | HTBE |
| *ATF3* | 467 | 1.52996584 | 4.77239E-06 | HTBE |
| *ATP10A* | 57194 | 1.658247655 | 4.18906E-35 | A549 |
| *ATP10A* | 57194 | 4.849310574 | 2.66776E-62 | HTBE |
| *ATP10D* | 57205 | 2.405236073 | 9.08593E-44 | A549 |
| *B2M* | 567 | 2.560006927 | 0 | A549 |
| *B3GALNT1* | 8706 | 1.504957537 | 6.96522E-12 | HTBE |
| *B3GNT7* | 93010 | 1.790411364 | 2.28704E-26 | HTBE |
| *BATF2* | 116071 | 5.397707298 | 2.51415E-66 | A549 |
| *BATF2* | 116071 | 5.786143995 | 1.2395E-149 | HTBE |
| *BCL2A1* | 597 | 2.74254995 | 1.45769E-06 | A549 |
| *BCL2L14* | 79370 | 5.965750663 | 0.000646764 | HTBE |
| *BIRC3* | 330 | 2.351798056 | 2.4847E-188 | A549 |
| *BISPR* | 105221694 | 5.607614508 | 0.001644859 | A549 |
| *BISPR* | 105221694 | 4.871273377 | 1.05495E-08 | HTBE |
| *BLNK* | 29760 | 2.482037808 | 0.002345333 | HTBE |
| *BMP4* | 652 | 2.166448949 | 0.012603622 | HTBE |
| *BST2* | 684 | 8.23636146 | 1.35954E-40 | A549 |
| *BST2* | 684 | 4.428403621 | 9.3017E-106 | HTBE |

| | | | | |
|---|---|---|---|---|
| *BTC* | 685 | 1.778000482 | 2.30181E-07 | A549 |
| *BTC* | 685 | 4.556548566 | 2.34625E-11 | HTBE |
| *BTN3A1* | 11119 | 1.886959954 | 4.01447E-30 | A549 |
| *BTN3A1* | 11119 | 2.168469407 | 1.70623E-18 | HTBE |
| *BTN3A2* | 11118 | 1.764704855 | 7.19044E-25 | A549 |
| *BTN3A3* | 10384 | 1.864671535 | 1.21182E-10 | A549 |
| *BTN3A3* | 10384 | 2.493365791 | 1.00321E-18 | HTBE |
| *C11orf86* | 254439 | 3.139898134 | 1.01312E-13 | A549 |
| *C12orf74* | 440107 | 6.796865388 | 0.000691362 | HTBE |
| *C15orf48* | 84419 | 4.295184077 | 1.01671E-17 | A549 |
| *C17orf67* | 339210 | 2.016616026 | 0.000340812 | HTBE |
| *C19orf66* | 55337 | 2.936821609 | 2.3887E-99 | A549 |
| *C19orf66* | 55337 | 3.40440562 | 5.97E-137 | HTBE |
| *C1orf74* | 148304 | 1.59469431 | 4.28811E-28 | HTBE |
| *C1R* | 715 | 1.613775599 | 2.88144E-57 | A549 |
| *C1R* | 715 | 4.385111554 | 3.02301E-44 | HTBE |
| *C1S* | 716 | 4.498407138 | 6.65336E-20 | HTBE |
| *C3AR1* | 719 | 5.397195306 | 7.90412E-14 | HTBE |
| *C4orf33* | 132321 | 1.715069534 | 8.61196E-19 | A549 |
| *C4orf33* | 132321 | 1.823775911 | 9.08914E-13 | HTBE |
| *C5orf56* | 441108 | 2.51151632 | 3.15508E-05 | A549 |
| *C5orf56* | 441108 | 3.96380804 | 7.15022E-09 | HTBE |
| *C6orf141* | 135398 | 1.757899234 | 7.25337E-15 | A549 |
| *CAPNS2* | 84290 | 2.077938047 | 1.02948E-12 | HTBE |
| *CARD16* | 114769 | 3.393855673 | 0.047139356 | A549 |
| *CARD16* | 114769 | 3.011406249 | 8.6582E-07 | HTBE |
| *CASP1* | 834 | 4.892625967 | 4.4835E-11 | A549 |
| *CASP1* | 834 | 2.802274371 | 1.32287E-58 | HTBE |
| *CASP10* | 843 | 1.932446185 | 4.46073E-47 | HTBE |
| *CASP4* | 837 | 1.563505141 | 3.18246E-41 | HTBE |
| *CASP7* | 840 | 1.625803644 | 2.22524E-63 | A549 |
| *CASP7* | 840 | 2.037898468 | 1.1298E-64 | HTBE |
| *CBR3* | 874 | 2.236082279 | 1.75831E-22 | HTBE |
| *CCDC109B* | 55013 | 2.112865104 | 7.28982E-53 | HTBE |
| *CCL2* | 6347 | 1.867998405 | 2.08989E-15 | A549 |
| *CCNA1* | 8900 | 3.265325389 | 1.0603E-165 | HTBE |
| *CD274* | 29126 | 3.002137966 | 8.53526E-11 | A549 |
| *CD274* | 29126 | 1.503758151 | 8.41796E-29 | HTBE |
| *CD68* | 968 | 1.868230879 | 8.49355E-47 | HTBE |
| *CEACAM1* | 634 | 2.256682805 | 8.64086E-11 | A549 |
| *CEACAM1* | 634 | 2.548981338 | 1.27309E-28 | HTBE |
| *CEBPB* | 1051 | 1.668417487 | 4.6685E-27 | A549 |
| *CFB* | 629 | 2.017592847 | 7.70606E-36 | A549 |
| *CFB* | 629 | 2.540399981 | 5.64898E-46 | HTBE |
| *CHORDC1* | 26973 | 2.571522188 | 1.3885E-188 | A549 |
| *CHSY3* | 337876 | 3.371159187 | 0.000489361 | HTBE |
| *CLDN23* | 137075 | 2.454362467 | 0.000293827 | HTBE |
| *CLEC2B* | 9976 | 2.018654905 | 0.02872862 | HTBE |

| | | | | |
|---|---|---|---|---|
| *CLEC7A* | 64581 | 4.406417851 | 0.03471953 | A549 |
| *CMPK2* | 129607 | 7.525087319 | 3.6256E-253 | A549 |
| *CMPK2* | 129607 | 13.14962244 | 2.67816E-19 | HTBE |
| *CMTR1* | 23070 | 2.021695482 | 1.85817E-86 | HTBE |
| *CNP* | 1267 | 2.033805269 | 3.6829E-103 | HTBE |
| *CTB-178M22.2* | | 2.884992187 | 0.003113938 | A549 |
| *CTSS* | 1520 | 1.659601974 | 3.77074E-20 | HTBE |
| *CX3CL1* | 6376 | 3.185675187 | 6.52834E-34 | HTBE |
| *CXCL10* | 3627 | 10.99629881 | 0 | HTBE |
| *CXCL11* | 6373 | 5.691227188 | 0.008456671 | A549 |
| *CXCL11* | 6373 | 9.818482198 | 6.395E-294 | HTBE |
| *CXCL17* | 284340 | 1.551063931 | 0.01408254 | HTBE |
| *CXCL9* | 4283 | 5.616661484 | 2.05675E-05 | HTBE |
| *CYP2J2* | 1573 | 3.483671072 | 9.48696E-07 | A549 |
| *CYP2J2* | 1573 | 5.261988763 | 9.20861E-38 | HTBE |
| *DAPP1* | 27071 | 1.58058548 | 1.27086E-37 | HTBE |
| *DAW1* | 164781 | 1.872259252 | 0.015875365 | A549 |
| *DDO* | 8528 | 5.355210768 | 2.53782E-05 | A549 |
| *DDX58* | 23586 | 5.21044185 | 0 | A549 |
| *DDX58* | 23586 | 5.582505267 | 0 | HTBE |
| *DDX60* | 55601 | 4.620112893 | 0 | A549 |
| *DDX60* | 55601 | 5.59249431 | 0 | HTBE |
| *DDX60L* | 91351 | 4.474753871 | 0 | A549 |
| *DDX60L* | 91351 | 5.277763948 | 0 | HTBE |
| *DHX58* | 79132 | 5.735470906 | 7.21413E-48 | A549 |
| *DHX58* | 79132 | 4.555061694 | 8.7886E-143 | HTBE |
| *DIO2* | 1734 | 2.390481279 | 8.88739E-74 | A549 |
| *DNAH17* | 8632 | 2.4441887 | 0.004548093 | A549 |
| *DNAJA1* | 3301 | 2.110201134 | 9.2695E-242 | A549 |
| *DNAJC6* | 9829 | 1.739536601 | 0.000141093 | HTBE |
| *DTX3L* | 151636 | 3.6695701 | 0 | A549 |
| *DTX3L* | 151636 | 3.457521629 | 2.9542E-259 | HTBE |
| *DUOX2* | 50506 | 4.298274026 | 5.93324E-29 | HTBE |
| *DUOXA2* | 405753 | 6.019817983 | 2.45801E-57 | HTBE |
| *EHD4* | 30844 | 1.522018711 | 2.89059E-40 | A549 |
| *EIF2AK2* | 5610 | 2.721895023 | 0 | A549 |
| *EIF2AK2* | 5610 | 3.473477612 | 1.98E-202 | HTBE |
| *ELOVL3* | 83401 | 3.003665275 | 0.004170747 | A549 |
| *EPSTI1* | 94240 | 6.765274225 | 1.38031E-21 | A549 |
| *EPSTI1* | 94240 | 6.096792301 | 5.5256E-229 | HTBE |
| *ERAP1* | 51752 | 1.574793896 | 8.91606E-57 | A549 |
| *ERAP2* | 64167 | 2.510722476 | 3.75291E-27 | A549 |
| *EREG* | 2069 | 2.007781355 | 2.6658E-100 | A549 |
| *ETV7* | 51513 | 5.709443928 | 2.20979E-11 | A549 |
| *ETV7* | 51513 | 6.16450027 | 1.67504E-82 | HTBE |
| *EVA1A* | 84141 | 1.565896263 | 0.001123882 | A549 |
| *EXOC3L1* | 283849 | 5.513969285 | 0.002201836 | A549 |
| *EXOC3L1* | 283849 | 6.747348092 | 0.000105333 | HTBE |

| | | | | |
|---|---|---|---|---|
| *FAM122C* | 159091 | 4.286410549 | 6.78277E-06 | HTBE |
| *FAM46A* | 55603 | 3.427080776 | 4.88863E-43 | HTBE |
| *FAM46C* | 54855 | 4.229368295 | 5.66898E-08 | HTBE |
| *FBXO6* | 26270 | 1.530071974 | 1.34839E-13 | A549 |
| *FBXO6* | 26270 | 2.812695348 | 2.34908E-64 | HTBE |
| *FGD2* | 221472 | 3.304979719 | 3.35068E-10 | HTBE |
| *FLT3LG* | 2323 | 2.428124808 | 0.000471883 | A549 |
| *FLT3LG* | 2323 | 2.055521032 | 0.007773529 | HTBE |
| *FSIP1* | 161835 | 2.032581401 | 0.007993015 | HTBE |
| *FYB* | 2533 | 1.572874201 | 1.80115E-21 | HTBE |
| *GALM* | 130589 | 3.742038867 | 1.39571E-21 | HTBE |
| *GBP1* | 2633 | 8.360270831 | 4.34577E-35 | A549 |
| *GBP1* | 2633 | 4.885426134 | 0 | HTBE |
| *GBP1P1* | 400759 | 3.032056929 | 4.20546E-07 | HTBE |
| *GBP2* | 2634 | 1.883750407 | 7.67831E-10 | HTBE |
| *GBP3* | 2635 | 3.646931788 | 2.33784E-91 | A549 |
| *GBP3* | 2635 | 2.356545436 | 3.6257E-67 | HTBE |
| *GBP4* | 115361 | 10.80744446 | 1.40585E-12 | A549 |
| *GBP4* | 115361 | 7.015764242 | 1.6616E-185 | HTBE |
| *GBP5* | 115362 | 6.016749709 | 0.002701253 | A549 |
| *GBP5* | 115362 | 6.087614054 | 3.3476E-99 | HTBE |
| *GCH1* | 2643 | 2.031421823 | 3.64231E-24 | HTBE |
| *GCNT4* | 51301 | 1.638388693 | 0.00188552 | A549 |
| *GCNT4* | 51301 | 1.601584196 | 2.03835E-06 | HTBE |
| *GIMAP2* | 26157 | 5.06600455 | 1.33575E-06 | A549 |
| *GIMAP2* | 26157 | 4.550060825 | 4.17319E-07 | HTBE |
| *GMPR* | 2766 | 3.849134782 | 4.90977E-08 | A549 |
| *GMPR* | 2766 | 8.089222495 | 3.31649E-32 | HTBE |
| *GNB4* | 59345 | 1.721328468 | 8.70639E-52 | HTBE |
| *GSDMD* | 79792 | 1.916109588 | 9.99899E-53 | HTBE |
| *HAPLN3* | 145864 | 2.516153494 | 4.91749E-17 | A549 |
| *HAS3* | 3038 | 1.719009183 | 2.75961E-11 | A549 |
| *HCP5* | 10866 | 1.605780968 | 0.015351215 | A549 |
| *HDX* | 139324 | 1.975895667 | 8.61107E-13 | A549 |
| *HDX* | 139324 | 1.920206035 | 7.89678E-13 | HTBE |
| *HELZ2* | 85441 | 3.604682944 | 7.3831E-283 | A549 |
| *HELZ2* | 85441 | 4.186556512 | 0 | HTBE |
| *HERC5* | 51191 | 5.130356757 | 0 | A549 |
| *HERC5* | 51191 | 9.50305277 | 6.11417E-72 | HTBE |
| *HERC6* | 55008 | 3.17336526 | 4.4827E-149 | A549 |
| *HERC6* | 55008 | 5.228416091 | 4.2062E-228 | HTBE |
| *HES4* | 57801 | 1.976785767 | 4.89406E-15 | HTBE |
| *HLA-B* | 3106 | 3.040675491 | 2.38453E-95 | A549 |
| *HLA-B* | 3106 | 2.797798802 | 3.7098E-187 | HTBE |
| *HLA-C* | 3107 | 1.823151727 | 1.66194E-61 | A549 |
| *HLA-C* | 3107 | 1.770912193 | 4.2853E-107 | HTBE |
| *HLA-E* | 3133 | 2.00883454 | 8.5585E-165 | A549 |
| *HLA-E* | 3133 | 1.591935188 | 1.67471E-85 | HTBE |

| | | | | |
|---|---|---|---|---|
| *HLA-F* | 3134 | 3.602008398 | 1.04741E-66 | A549 |
| *HLA-F* | 3134 | 3.735342024 | 2.5367E-131 | HTBE |
| *HRASLS2* | 54979 | 3.115214842 | 2.06318E-08 | A549 |
| *HS3ST1* | 9957 | 1.533582905 | 8.28557E-27 | HTBE |
| *HSH2D* | 84941 | 1.621572636 | 0.00021747 | A549 |
| *HSH2D* | 84941 | 5.28258679 | 1.0187E-165 | HTBE |
| *HSP90AA1* | 3320 | 1.725291739 | 9.79208E-17 | A549 |
| *HSPA1A* | 3303 | 2.417505007 | 2.0296E-176 | A549 |
| *HSPA1B* | 3304 | 2.309362355 | 1.8569E-125 | A549 |
| *HSPA4L* | 22824 | 1.514998421 | 1.96642E-33 | A549 |
| *HSPA8* | 3312 | 1.835040561 | 1.7057E-249 | A549 |
| *HSPH1* | 10808 | 1.806389018 | 6.3851E-117 | A549 |
| *ID1* | 3397 | 2.399967262 | 1.2342E-258 | A549 |
| *ID2* | 3398 | 2.225243692 | 5.4059E-148 | A549 |
| *ID3* | 3399 | 2.272167265 | 1.1425E-113 | A549 |
| *ID4* | 3400 | 4.232581665 | 2.19408E-47 | A549 |
| *IDO1* | 3620 | 9.199012738 | 9.36498E-18 | A549 |
| *IDO1* | 3620 | 5.489690813 | 5.40952E-79 | HTBE |
| *IFI16* | 3428 | 5.240392019 | 3.3929E-155 | A549 |
| *IFI16* | 3428 | 2.677018697 | 1.4466E-176 | HTBE |
| *IFI27* | 3429 | 8.901304644 | 3.45713E-87 | A549 |
| *IFI27* | 3429 | 3.528053588 | 1.5341E-277 | HTBE |
| *IFI30* | 10437 | 1.767895466 | 2.53811E-30 | A549 |
| *IFI30* | 10437 | 1.80491655 | 9.45847E-20 | HTBE |
| *IFI35* | 3430 | 3.197296442 | 0 | A549 |
| *IFI35* | 3430 | 5.308570226 | 1.712E-261 | HTBE |
| *IFI44* | 10561 | 7.960093397 | 5.27056E-74 | A549 |
| *IFI44* | 10561 | 7.346983028 | 8.4112E-194 | HTBE |
| *IFI44L* | 10964 | 11.6594713 | 1.33033E-14 | A549 |
| *IFI44L* | 10964 | 10.63764547 | 1.33326E-57 | HTBE |
| *IFI6* | 2537 | 7.286546032 | 0 | A549 |
| *IFI6* | 2537 | 7.011007713 | 0 | HTBE |
| *IFIH1* | 64135 | 6.542085348 | 0 | A549 |
| *IFIH1* | 64135 | 6.057909581 | 0 | HTBE |
| *IFIT1* | 3434 | 9.187634915 | 0 | A549 |
| *IFIT1* | 3434 | 10.98666065 | 0 | HTBE |
| *IFIT2* | 3433 | 4.847912717 | 0 | A549 |
| *IFIT2* | 3433 | 9.097822455 | 0 | HTBE |
| *IFIT3* | 3437 | 6.818982917 | 0 | A549 |
| *IFIT3* | 3437 | 8.573467711 | 2.2359E-132 | HTBE |
| *IFIT5* | 24138 | 3.092454355 | 1.7837E-280 | A549 |
| *IFIT5* | 24138 | 4.198114971 | 6.0656E-206 | HTBE |
| *IFITM1* | 8519 | 5.810286868 | 1.57745E-97 | A549 |
| *IFITM1* | 8519 | 5.121351399 | 0 | HTBE |
| *IFITM2* | 10581 | 1.882075432 | 4.38471E-29 | HTBE |
| *IFITM3* | 10410 | 3.493919777 | 1.27328E-80 | A549 |
| *IFITM3* | 10410 | 1.836916315 | 2.09852E-79 | HTBE |
| *IL11* | 3589 | 1.705324835 | 1.99152E-05 | A549 |

| | | | | |
|---|---|---|---|---|
| *IL12A* | 3592 | 3.287341804 | 0.000197311 | A549 |
| *IL15* | 3600 | 1.944367643 | 1.2247E-08 | HTBE |
| *IL15RA* | 3601 | 1.987550379 | 5.52457E-07 | A549 |
| *IL15RA* | 3601 | 3.02626251 | 2.16462E-42 | HTBE |
| *IL22RA1* | 58985 | 2.425449381 | 1.7463E-16 | HTBE |
| *IL4I1* | 259307 | 3.403951685 | 0.013650844 | A549 |
| *IL4I1* | 259307 | 3.374216035 | 0.013783113 | HTBE |
| *IL7* | 3574 | 7.056442776 | 6.38532E-05 | A549 |
| *IL7* | 3574 | 3.751266264 | 3.86416E-06 | HTBE |
| *IRF1* | 3659 | 2.537174574 | 2.989E-107 | HTBE |
| *IRF7* | 3665 | 4.951144734 | 8.2166E-286 | A549 |
| *IRF7* | 3665 | 5.090719375 | 0 | HTBE |
| *IRF9* | 10379 | 3.155455968 | 2.298E-173 | A549 |
| *IRF9* | 10379 | 1.937721081 | 4.96235E-61 | HTBE |
| *ISG15* | 9636 | 7.544119651 | 0 | A549 |
| *ISG15* | 9636 | 7.147989086 | 0 | HTBE |
| *ISG20* | 3669 | 2.704127006 | 3.8175E-127 | A549 |
| *ISG20* | 3669 | 1.899899163 | 2.04872E-83 | HTBE |
| *ITGB6* | 3694 | 2.225938576 | 8.82765E-05 | A549 |
| *JAG1* | 182 | 1.713021582 | 6.3025E-124 | A549 |
| *JAK2* | 3717 | 1.739181691 | 4.16476E-24 | HTBE |
| *KBTBD8* | 84541 | 2.074797881 | 8.7571E-05 | A549 |
| *KBTBD8* | 84541 | 1.593756103 | 5.27266E-08 | HTBE |
| *KCTD21-AS1* | 100289388 | 1.625691267 | 0.030358071 | A549 |
| *KIAA1644* | 85352 | 1.747379541 | 0.020667722 | A549 |
| *KLB* | 152831 | 1.903036703 | 0.032023888 | A549 |
| *KRT17* | 3872 | 3.716402205 | 0.000166231 | A549 |
| *LAMP3* | 27074 | 7.042765595 | 3.23686E-19 | A549 |
| *LAMP3* | 27074 | 6.833618863 | 3.09178E-79 | HTBE |
| *LAP3* | 51056 | 2.726921109 | 0 | A549 |
| *LAP3* | 51056 | 2.778604731 | 2.0286E-131 | HTBE |
| *LBP* | 3929 | 2.571853599 | 0.045729174 | A549 |
| *LGALS17A* | 400696 | 6.071393578 | 0.002381765 | A549 |
| *LGALS9* | 3965 | 4.524851159 | 4.58187E-19 | A549 |
| *LGALS9* | 3965 | 5.722672829 | 4.36959E-45 | HTBE |
| *LGMN* | 5641 | 1.512156181 | 7.6388E-37 | HTBE |
| *LIFR* | 3977 | 1.658154193 | 2.99169E-11 | HTBE |
| *LMO2* | 4005 | 2.803987391 | 3.87585E-06 | A549 |
| *LMO2* | 4005 | 4.680566734 | 1.25681E-39 | HTBE |
| *LNX1* | 84708 | 1.594551407 | 0.027051477 | HTBE |
| *LOC100419583* | 100419583 | 3.215667183 | 2.4537E-114 | A549 |
| *LOC100419583* | 100419583 | 3.93946753 | 5.7758E-141 | HTBE |
| *LOC101927027* | 101927027 | 1.585843716 | 2.76648E-08 | HTBE |
| *LOC101927746* | 101927746 | 1.572904341 | 4.72584E-09 | A549 |
| *LOC101929723* | 101929723 | 1.725244456 | 0.002400174 | A549 |
| *LOC102577426* | 102577426 | 2.200071793 | 0.022968769 | A549 |
| *LOC102724163* | 102724163 | 7.704876659 | 1.78323E-05 | HTBE |
| *LOC152225* | 152225 | 1.809812679 | 0.000102682 | HTBE |

| | | | | |
|---|---|---|---|---|
| *LRP2* | 4036 | 8.230159641 | 1.74751E-06 | HTBE |
| *LUCAT1* | 100505994 | 2.414464256 | 0.030843571 | HTBE |
| *LY6E* | 4061 | 1.917399069 | 8.72298E-84 | HTBE |
| *LYAR* | 55646 | 1.506182734 | 4.1923E-29 | A549 |
| *LYSMD2* | 256586 | 2.381458717 | 2.63548E-28 | HTBE |
| *MAK* | 4117 | 6.360023599 | 0.000422248 | HTBE |
| *MAN1A1* | 4121 | 1.55948907 | 2.26863E-05 | HTBE |
| *MAP2* | 4133 | 1.775526542 | 1.92786E-10 | A549 |
| *MASTL* | 84930 | 2.000341814 | 2.9366E-41 | HTBE |
| *MB21D2* | 151963 | 1.811299342 | 8.64665E-24 | A549 |
| *MCTP1* | 79772 | 1.612166106 | 2.99853E-15 | A549 |
| *MIR3193* | 100422904 | 1.922052873 | 0.001248847 | A549 |
| *MLKL* | 197259 | 2.28557198 | 6.21373E-82 | A549 |
| *MLKL* | 197259 | 2.290190993 | 3.35867E-63 | HTBE |
| *MMP13* | 4322 | 3.456673215 | 0.042252479 | A549 |
| *MMP13* | 4322 | 5.858803855 | 1.3113E-108 | HTBE |
| *MX1* | 4599 | 9.430581588 | 0 | A549 |
| *MX1* | 4599 | 9.156963208 | 0 | HTBE |
| *MX2* | 4600 | 11.40154446 | 3.05088E-14 | A549 |
| *MX2* | 4600 | 11.73198362 | 1.61208E-70 | HTBE |
| *MYD88* | 4615 | 2.187607406 | 1.78036E-89 | A549 |
| *MYD88* | 4615 | 2.332122943 | 1.59579E-77 | HTBE |
| *NAPSB* | 256236 | 6.300643705 | 0.001080589 | A549 |
| *NAT8* | 9027 | 2.993231899 | 0.046507451 | A549 |
| *NCOA7* | 135112 | 1.65412049 | 3.6885E-134 | A549 |
| *NCOA7* | 135112 | 3.049976985 | 1.11238E-94 | HTBE |
| *NEURL3* | 93082 | 3.629546758 | 0.000308205 | HTBE |
| *NFASC* | 23114 | 2.120901101 | 0.003489641 | HTBE |
| *NFE2L3* | 9603 | 2.678939963 | 9.27066E-91 | A549 |
| *NFE2L3* | 9603 | 1.687764829 | 4.09786E-27 | HTBE |
| *NID2* | 22795 | 1.509923761 | 0.016556005 | A549 |
| *NLRC5* | 84166 | 3.359841897 | 7.5263E-107 | A549 |
| *NLRC5* | 84166 | 5.257527402 | 2.849E-133 | HTBE |
| *NMI* | 9111 | 3.403664332 | 1.2169E-213 | A549 |
| *NMI* | 9111 | 4.015540032 | 5.67215E-88 | HTBE |
| *NOC3L* | 64318 | 1.61556468 | 3.41388E-70 | A549 |
| *NOD2* | 64127 | 2.790602819 | 1.93672E-05 | HTBE |
| *NOS2* | 4843 | 2.723204803 | 0.00343419 | A549 |
| *NRIR* | 104326052 | 6.451769503 | 0.000622781 | A549 |
| *NT5C3A* | 51251 | 2.971493349 | 0 | A549 |
| *NT5C3A* | 51251 | 3.691098797 | 1.9039E-136 | HTBE |
| *NUB1* | 51667 | 2.35009897 | 1.09909E-74 | HTBE |
| *NUDCD1* | 84955 | 1.501447274 | 4.14192E-73 | A549 |
| *NUDCD1* | 84955 | 1.864520991 | 4.04799E-56 | HTBE |
| *NUPR1* | 26471 | 4.423079184 | 5.7401E-199 | HTBE |
| *OAS1* | 4938 | 2.675496466 | 0 | A549 |
| *OAS1* | 4938 | 7.969619553 | 0 | HTBE |
| *OAS2* | 4939 | 9.858507948 | 4.0361E-118 | A549 |

| | | | | |
|---|---|---|---|---|
| *OAS2* | 4939 | 5.851984734 | 5.668E-172 | HTBE |
| *OAS3* | 4940 | 3.009949927 | 0 | A549 |
| *OAS3* | 4940 | 4.566071549 | 0 | HTBE |
| *OASL* | 8638 | 7.590845226 | 0 | A549 |
| *OASL* | 8638 | 9.786620504 | 3.8634E-186 | HTBE |
| *ODF3B* | 440836 | 1.563478497 | 7.29334E-07 | A549 |
| *ODF3B* | 440836 | 2.33397746 | 2.00697E-11 | HTBE |
| *OGFR* | 11054 | 1.709721445 | 4.97469E-94 | A549 |
| *OGFR* | 11054 | 2.032167344 | 1.44852E-63 | HTBE |
| *OVOL1* | 5017 | 1.958317523 | 6.40432E-07 | HTBE |
| *PARP10* | 84875 | 6.291980347 | 4.0712E-219 | A549 |
| *PARP10* | 84875 | 3.481683215 | 1.3584E-164 | HTBE |
| *PARP12* | 64761 | 3.7820442 | 0 | A549 |
| *PARP12* | 64761 | 3.49588892 | 4.9678E-258 | HTBE |
| *PARP14* | 54625 | 4.226158392 | 0 | A549 |
| *PARP14* | 54625 | 4.807698849 | 0 | HTBE |
| *PARP9* | 83666 | 5.02582621 | 0 | A549 |
| *PARP9* | 83666 | 4.707153144 | 0 | HTBE |
| *PATL2* | 197135 | 4.784271372 | 1.0623E-05 | A549 |
| *PCDH17* | 27253 | 5.312391016 | 0.00010289 | HTBE |
| *PCDH9* | 5101 | 1.985042162 | 4.69764E-05 | HTBE |
| *PCDHAC2* | 56134 | 1.560711588 | 9.13209E-05 | HTBE |
| *PCGF5* | 84333 | 2.194699623 | 4.69567E-45 | HTBE |
| *PDCD1LG2* | 80380 | 4.586643996 | 0.023973973 | A549 |
| *PDCD1LG2* | 80380 | 2.119729944 | 1.4754E-19 | HTBE |
| *PDGFRL* | 5157 | 1.979894497 | 4.11483E-38 | A549 |
| *PDZD2* | 23037 | 4.570445305 | 0.023915208 | A549 |
| *PDZD2* | 23037 | 2.472990229 | 2.00469E-30 | HTBE |
| *PHF11* | 51131 | 2.358312433 | 6.03966E-35 | A549 |
| *PHF11* | 51131 | 2.851815709 | 1.5133E-61 | HTBE |
| *PIK3AP1* | 118788 | 4.038014397 | 4.78156E-47 | A549 |
| *PITRM1-AS1* | 100507034 | 1.97646374 | 0.037717373 | A549 |
| *PKIB* | 5570 | 1.52719833 | 0.000991347 | HTBE |
| *PLAC8* | 51316 | 3.121176542 | 1.68917E-05 | HTBE |
| *PLEKHA4* | 57664 | 5.202560945 | 1.17087E-46 | A549 |
| *PLEKHA4* | 57664 | 3.854315583 | 6.2207E-112 | HTBE |
| *PLEKHG7* | 440107 | 2.923827534 | 0.041590964 | HTBE |
| *PLSCR1* | 5359 | 2.901998696 | 0 | A549 |
| *PLSCR1* | 5359 | 3.440218906 | 3.0405E-245 | HTBE |
| *PLSCR4* | 57088 | 2.733491344 | 0.00384407 | HTBE |
| *PLXNC1* | 10154 | 2.033322585 | 0.044141884 | HTBE |
| *PMAIP1* | 5366 | 1.926641262 | 1.61199E-31 | HTBE |
| *PML* | 5371 | 2.855376923 | 2.43857E-53 | A549 |
| *PML* | 5371 | 2.309771362 | 8.05803E-97 | HTBE |
| *PNPT1* | 87178 | 3.121005362 | 0 | A549 |
| *PNPT1* | 87178 | 3.243928222 | 5.2827E-224 | HTBE |
| *POMGNT2* | 84892 | 1.501967417 | 5.83241E-11 | HTBE |
| *PPM1K* | 152926 | 2.317199219 | 2.35977E-20 | A549 |

| | | | | |
|---|---|---|---|---|
| *PPM1K* | 152926 | 3.370840662 | 1.1375E-101 | HTBE |
| *PRICKLE4* | 29964 | 1.623061252 | 9.61133E-05 | HTBE |
| *PRR15* | 222171 | 2.692996576 | 2.0025E-07 | HTBE |
| *PSMB8* | 5696 | 3.412794383 | 2.2953E-235 | A549 |
| *PSMB8* | 5696 | 2.217237391 | 5.4224E-109 | HTBE |
| *PSMB8-AS1* | 100507463 | 3.872024935 | 4.56721E-29 | A549 |
| *PSMB8-AS1* | 100507463 | 3.821680179 | 8.46002E-10 | HTBE |
| *PSMB9* | 5698 | 4.664414674 | 9.3562E-215 | A549 |
| *PSMB9* | 5698 | 3.249785442 | 3.7709E-125 | HTBE |
| *PSME2* | 5721 | 1.924559083 | 3.3031E-138 | A549 |
| *PTCH2* | 8643 | 1.640682095 | 0.00177911 | A549 |
| *PVRL4* | 81607 | 4.560733668 | 0.024700492 | A549 |
| *RAB19* | 401409 | 3.47557291 | 0.010255979 | HTBE |
| *RAB39A* | 54734 | 1.706193855 | 0.000962411 | A549 |
| *RARRES3* | 5920 | 3.299923209 | 7.35121E-69 | A549 |
| *RASGRP3* | 25780 | 2.611646438 | 6.38161E-30 | A549 |
| *RASGRP3* | 25780 | 7.011191462 | 3.86455E-27 | HTBE |
| *RBCK1* | 10616 | 1.811921917 | 7.58602E-54 | HTBE |
| *RBM11* | 54033 | 5.063806645 | 8.54607E-08 | HTBE |
| *RBM43* | 375287 | 2.570120973 | 4.44196E-10 | HTBE |
| *REC8* | 9985 | 2.380259586 | 2.48708E-09 | A549 |
| *RET* | 5979 | 2.100162511 | 0.001767566 | A549 |
| *RET* | 5979 | 3.087275639 | 6.6352E-06 | HTBE |
| *RGS22* | 26166 | 3.693395153 | 0.000193148 | A549 |
| *RIMS2* | 9699 | 2.716765294 | 4.08307E-40 | HTBE |
| *RN7SL2* | 378706 | 1.518199056 | 1.37575E-29 | A549 |
| *RNA18S5* | 110255169 | 1.641538473 | 0.024920703 | A549 |
| *RNF114* | 55905 | 1.7462248 | 8.15304E-58 | HTBE |
| *RNF144A* | 9781 | 2.693818428 | 0.013781924 | HTBE |
| *RNF213* | 57674 | 2.481267463 | 0 | A549 |
| *RNF213* | 57674 | 3.282401156 | 5.47752E-36 | HTBE |
| *RSAD2* | 91543 | 7.618217962 | 4.27988E-45 | A549 |
| *RSAD2* | 91543 | 11.76158519 | 1.4345E-125 | HTBE |
| *RTKN2* | 219790 | 1.509140398 | 0.004645668 | HTBE |
| *RTP4* | 64108 | 8.600345571 | 8.20614E-08 | A549 |
| *RTP4* | 64108 | 5.902256113 | 4.02768E-57 | HTBE |
| *RUFY4* | 285180 | 6.835906039 | 2.87722E-05 | A549 |
| *SAA2* | 6288 | 1.733446994 | 8.07826E-05 | HTBE |
| *SAMD9* | 54809 | 4.458936496 | 0 | A549 |
| *SAMD9* | 54809 | 4.27882783 | 0 | HTBE |
| *SAMD9L* | 219285 | 5.656938876 | 4.7089E-292 | A549 |
| *SAMD9L* | 219285 | 6.428648945 | 8.3468E-215 | HTBE |
| *SAMHD1* | 25939 | 3.208708588 | 2.4849E-276 | A549 |
| *SAMHD1* | 25939 | 4.069743783 | 1.6584E-260 | HTBE |
| *SCLT1* | 132320 | 1.515360017 | 7.58853E-24 | A549 |
| *SDPR* | 8436 | 1.998101844 | 8.51225E-07 | HTBE |
| *SEC16B* | 89866 | 1.515643788 | 5.23408E-10 | A549 |
| *SECTM1* | 6398 | 3.062310099 | 1.77543E-06 | A549 |

| | | | | |
|---|---|---|---|---|
| *SECTM1* | 6398 | 6.220006181 | 1.67766E-59 | HTBE |
| *SEMA3D* | 223117 | 4.463677164 | 0.00557499 | HTBE |
| *SERPINE1* | 5054 | 1.924488732 | 1.8812E-119 | A549 |
| *SERPING1* | 710 | 4.708746652 | 0.01784781 | A549 |
| *SIDT1* | 54847 | 3.964238025 | 2.65374E-21 | HTBE |
| *SIRPB2* | 284759 | 2.213229476 | 4.36731E-06 | HTBE |
| *SKIDA1* | 387640 | 3.268405049 | 0.010242075 | HTBE |
| *SLC15A3* | 51296 | 9.366673072 | 2.4078E-09 | A549 |
| *SLC15A3* | 51296 | 5.718874911 | 2.63661E-37 | HTBE |
| *SLC16A4* | 9122 | 1.626337649 | 7.4577E-22 | HTBE |
| *SLC25A28* | 81894 | 2.364519945 | 1.26553E-44 | HTBE |
| *SLC2A12* | 154091 | 2.211108522 | 2.72239E-15 | HTBE |
| *SLC6A14* | 11254 | 1.745160098 | 2.30669E-42 | HTBE |
| *SLFN5* | 162394 | 1.761742048 | 2.31031E-86 | HTBE |
| *SNAI1* | 6615 | 2.149513588 | 0.004678953 | A549 |
| *SNORA14A* | 677801 | 2.628894084 | 0.033870923 | A549 |
| *SNORA22* | 677807 | 1.869425368 | 0.042598297 | A549 |
| *SNORA32* | 692063 | 1.534267418 | 1.3831E-05 | A549 |
| *SNORA53* | 677832 | 1.959631766 | 6.17867E-32 | A549 |
| *SNORA61* | 677838 | 1.535337193 | 0.000424382 | A549 |
| *SNORA65* | 26783 | 1.595950996 | 0.000624358 | A549 |
| *SNORD119* | 100113378 | 1.920846926 | 0.019057283 | A549 |
| *SNORD14C* | 85389 | 1.645065862 | 0.042537155 | A549 |
| *SNORD14D* | 85390 | 1.688548139 | 0.014291938 | A549 |
| *SNORD14E* | 85391 | 2.388594349 | 3.6948E-05 | A549 |
| *SNORD2* | 619567 | 1.505241426 | 2.53509E-05 | A549 |
| *SNORD47* | 26802 | 2.026003384 | 0.011242004 | A549 |
| *SNORD50B* | 692088 | 1.557567945 | 0.012898607 | A549 |
| *SNORD58A* | 26791 | 1.864434343 | 0.008906078 | A549 |
| *SNORD76* | 692196 | 1.748856971 | 0.001247406 | A549 |
| *SNORD78* | 692198 | 1.559559675 | 0.009720015 | A549 |
| *SNORD83A* | 116937 | 1.869389891 | 0.000640433 | A549 |
| *SNORD93* | 692210 | 1.955535794 | 0.010956635 | A549 |
| *SNORD99* | 692212 | 1.521488251 | 0.046547876 | A549 |
| *SOBP* | 55084 | 1.744707838 | 5.03119E-07 | HTBE |
| *SOCS1* | 8651 | 2.402218594 | 1.22693E-07 | HTBE |
| *SP100* | 6672 | 2.926020248 | 4.7768E-303 | A549 |
| *SP100* | 6672 | 3.246062971 | 1.1066E-241 | HTBE |
| *SP110* | 3431 | 3.859596628 | 1.5772E-120 | A549 |
| *SP110* | 3431 | 5.015360587 | 2.1516E-257 | HTBE |
| *SP140L* | 93349 | 3.828613164 | 3.08872E-75 | A549 |
| *SP140L* | 93349 | 3.665902401 | 1.61042E-78 | HTBE |
| *SPATA13* | 221178 | 1.568614459 | 1.77654E-07 | HTBE |
| *SPINK6* | 404203 | 2.186950924 | 0.042112361 | A549 |
| *SPTBN5* | 51332 | 1.877116623 | 0.03127691 | A549 |
| *ST8SIA4* | 7903 | 2.253721026 | 0.012409449 | HTBE |
| *STARD5* | 80765 | 3.531730082 | 3.46902E-86 | HTBE |
| *STAT1* | 6772 | 3.815563397 | 0 | A549 |

| | | | | |
|---|---|---|---|---|
| *STAT1* | 6772 | 4.492377453 | 0 | HTBE |
| *STAT2* | 6773 | 2.798046228 | 9.5286E-276 | A549 |
| *STAT2* | 6773 | 3.073934606 | 4.1808E-249 | HTBE |
| *STAT4* | 6775 | 1.854437757 | 1.75181E-20 | A549 |
| *SUSD3* | 203328 | 4.25047138 | 5.52966E-07 | HTBE |
| *SUSD4* | 55061 | 2.501241745 | 0.00010125 | HTBE |
| *SYNPO2* | 171024 | 3.466361054 | 3.38272E-08 | HTBE |
| *TAGAP* | 117289 | 3.768923044 | 0.002380757 | HTBE |
| *TAP1* | 6890 | 4.064002691 | 0 | A549 |
| *TAP1* | 6890 | 3.468549864 | 0 | HTBE |
| *TAP2* | 6891 | 2.755284245 | 2.1425E-158 | A549 |
| *TAP2* | 6891 | 1.799316076 | 6.45308E-61 | HTBE |
| *TCF4* | 6925 | 1.79728218 | 2.39784E-19 | HTBE |
| *TDRD7* | 23424 | 3.31755652 | 0 | A549 |
| *TDRD7* | 23424 | 4.196100992 | 1.2132E-253 | HTBE |
| *TFRC* | 7037 | 1.652985377 | 9.4196E-115 | A549 |
| *TGM2* | 7052 | 1.993868174 | 2.422E-193 | A549 |
| *THEMIS2* | 9473 | 4.975562193 | 2.24773E-32 | A549 |
| *THEMIS2* | 9473 | 4.441536895 | 1.0763E-186 | HTBE |
| *TIMP2* | 7077 | 1.513023513 | 1.55713E-07 | HTBE |
| *TLR2* | 7097 | 2.531325789 | 1.83357E-30 | HTBE |
| *TLR3* | 7098 | 4.052010659 | 1.48779E-61 | A549 |
| *TLR3* | 7098 | 4.309040935 | 9.08957E-67 | HTBE |
| *TMEM106A* | 113277 | 2.046080591 | 1.83672E-08 | HTBE |
| *TMEM140* | 55281 | 3.461434615 | 5.79592E-29 | A549 |
| *TMEM140* | 55281 | 8.875836123 | 7.0015E-24 | HTBE |
| *TMEM171* | 134285 | 1.652916324 | 2.1804E-07 | HTBE |
| *TMEM229B* | 161145 | 2.837695725 | 0.00038036 | A549 |
| *TMEM229B* | 161145 | 3.070315664 | 8.68435E-43 | HTBE |
| *TMEM27* | 57393 | 1.700029171 | 4.08702E-11 | A549 |
| *TMEM27* | 57393 | 3.960020805 | 5.37426E-10 | HTBE |
| *TMEM62* | 80021 | 1.864084633 | 7.58688E-40 | HTBE |
| *TNFRSF1B* | 7133 | 3.555838651 | 2.18483E-05 | A549 |
| *TNFSF10* | 8743 | 3.42398856 | 2.37069E-60 | A549 |
| *TNFSF10* | 8743 | 3.726875158 | 0 | HTBE |
| *TNFSF13* | 8741 | 1.80766552 | 0.004628957 | HTBE |
| *TNFSF13B* | 10673 | 6.154409781 | 7.0251E-14 | A549 |
| *TNFSF13B* | 10673 | 10.79054235 | 5.41643E-12 | HTBE |
| *TOR1B* | 27348 | 1.677575767 | 5.55695E-28 | HTBE |
| *TRANK1* | 9881 | 4.605778847 | 0 | A549 |
| *TRANK1* | 9881 | 4.663339091 | 8.2198E-181 | HTBE |
| *TREX1* | 11277 | 2.597418167 | 2.93785E-34 | HTBE |
| *TRIM14* | 9830 | 3.159652 | 7.0596E-113 | A549 |
| *TRIM14* | 9830 | 2.931610114 | 3.45945E-86 | HTBE |
| *TRIM21* | 6737 | 3.246198046 | 2.2277E-229 | A549 |
| *TRIM21* | 6737 | 2.188131871 | 1.05178E-93 | HTBE |
| *TRIM22* | 10346 | 7.579706335 | 7.0338E-141 | A549 |
| *TRIM22* | 10346 | 3.86329223 | 0 | HTBE |

| | | | | |
|---|---|---|---|---|
| *TRIM25* | 7706 | 2.594335866 | 1.0402E-255 | A549 |
| *TRIM25* | 7706 | 2.558673335 | 2.118E-173 | HTBE |
| *TRIM38* | 10475 | 1.548380595 | 1.26741E-40 | A549 |
| *TRIM38* | 10475 | 2.038056458 | 2.36213E-60 | HTBE |
| *TRIM5* | 85363 | 2.009465549 | 7.41539E-81 | A549 |
| *TRIM5* | 85363 | 2.439516224 | 2.1829E-108 | HTBE |
| *TRIM56* | 81844 | 2.132253154 | 7.29753E-36 | HTBE |
| *TRIM69* | 140691 | 2.99803859 | 0.024431198 | HTBE |
| *TRIML2* | 205860 | 2.001104982 | 3.19725E-80 | A549 |
| *TRPV2* | 51393 | 1.796194482 | 0.000464962 | HTBE |
| *TSPAN33* | 340348 | 2.103816162 | 0.00050057 | HTBE |
| *TTC39B* | 158219 | 1.509056879 | 3.09313E-10 | A549 |
| *TTLL11* | 158135 | 1.612460842 | 2.36685E-06 | HTBE |
| *TYMP* | 1890 | 3.058082929 | 9.84293E-89 | A549 |
| *TYMP* | 1890 | 2.357824783 | 1.241E-114 | HTBE |
| *UBA7* | 7318 | 5.644948969 | 9.57604E-47 | A549 |
| *UBA7* | 7318 | 2.050160081 | 5.22339E-44 | HTBE |
| *UBD* | 10537 | 3.666906896 | 1.83168E-08 | A549 |
| *UBE2L6* | 9246 | 4.79594311 | 0 | A549 |
| *UBE2L6* | 9246 | 3.731837478 | 4.9298E-299 | HTBE |
| *UNC93B1* | 81622 | 2.90600416 | 9.23919E-79 | HTBE |
| *USP18* | 11274 | 4.482760216 | 0 | A549 |
| *USP18* | 11274 | 5.765421572 | 0 | HTBE |
| *USP30-AS1* | 100131733 | 3.874738892 | 0.002249994 | A549 |
| *VTRNA1-3* | 56662 | 2.37064012 | 1.93792E-29 | A549 |
| *WARS* | 7453 | 2.197413367 | 1.4885E-127 | HTBE |
| *XAF1* | 54739 | 10.76344693 | 4.6047E-36 | A549 |
| *XAF1* | 54739 | 5.71079147 | 7.7692E-117 | HTBE |
| *XDH* | 7498 | 1.527108899 | 1.82293E-09 | A549 |
| *XRN1* | 54464 | 2.27799544 | 7.77975E-98 | HTBE |
| *ZBP1* | 81030 | 7.304043019 | 2.39271E-05 | A549 |
| *ZBP1* | 81030 | 9.471565511 | 6.23558E-10 | HTBE |
| *ZBTB42* | 100128927 | 2.193465778 | 2.83096E-16 | HTBE |
| *ZFP82* | 284406 | 1.882655762 | 3.79485E-08 | A549 |
| *ZFPM2* | 23414 | 3.166611029 | 1.74256E-06 | HTBE |
| *ZNF107* | 51427 | 1.687061171 | 1.81466E-07 | HTBE |
| *ZNF233* | 353355 | 1.574638547 | 0.008457291 | A549 |
| *ZNF284* | 342909 | 1.503283807 | 0.015507984 | A549 |
| *ZNF542P* | 147947 | 1.565857764 | 0.006572693 | A549 |
| *ZNFX1* | 57169 | 1.768463461 | 1.5281E-150 | A549 |
| *ZNFX1* | 57169 | 2.598352609 | 4.0425E-138 | HTBE |
| *ZPLD1* | 131368 | 1.54751312 | 0.036901896 | HTBE |

**Table S1 | Available ISGs**

cDNAs that were available as validated, full-sequence length clones (ORFeome library), from ISGs

| Gene Symbol | Gene ID |
|---|---|
| *ABLIM3* | 22885 |
| *ACSL5* | 51703 |
| *ACY3* | 91703 |
| *AHNAK2* | 113146 |
| *AIM2* | 9447 |
| *AKT3* | 10000 |
| *ALDH1A1* | 216 |
| *AMPH* | 273 |
| *ANGPTL1* | 9068 |
| *ANGPTL4* | 51129 |
| *ANKRD22* | 118932 |
| *APOBEC3A* | 200315 |
| *APOBEC3D* | 140564 |
| *APOBEC3F* | 200316 |
| *APOBEC3G* | 60489 |
| *APOL1* | 8542 |
| *APOL2* | 23780 |
| *APOL4* | 80832 |
| *APOL6* | 80830 |
| *AQP9* | 366 |
| *AREG* | 374 |
| *ARG2* | 384 |
| *ARHGAP17* | 55114 |
| *ARHGEF3* | 50650 |
| *ARNTL* | 406 |
| *ARRDC3* | 57561 |
| *ATF3* | 467 |
| *ATP10D* | 57205 |
| *AURKB* | 9212 |
| *B2M* | 567 |
| *B3GALNT1* | 8706 |
| *B4GALT5* | 9334 |
| *BAG1* | 573 |
| *BATF2* | 116071 |
| *BCL2L14* | 79370 |
| *BLNK* | 29760 |
| *BLVRA* | 644 |
| *BLZF1* | 8548 |
| *BMP4* | 652 |
| *BST2* | 684 |
| *BTC* | 685 |
| *BTN3A1* | 11119 |
| *BTN3A2* | 11118 |
| *BTN3A3* | 10384 |
| *C15orf48* | 84419 |

| | |
|---|---|
| *C17orf67* | 339210 |
| *C19orf66* | 55337 |
| *C1orf38* | 9473 |
| *C1R* | 715 |
| *C22orf28* | 51493 |
| *C3AR1* | 719 |
| *C3orf39* | 84892 |
| *C4orf33* | 132321 |
| *C5orf56* | 441108 |
| *C6orf141* | 135398 |
| *C9orf91* | 203197 |
| *CAPNS2* | 84290 |
| *CASP1* | 834 |
| *CASP10* | 843 |
| *CASP7* | 840 |
| *CBR3* | 874 |
| *CCDC109B* | 55013 |
| *CCDC75* | 253635 |
| *CCDC92* | 80212 |
| *CCL19* | 6363 |
| *CCL2* | 6347 |
| *CCL4* | 6351 |
| *CCL5* | 6352 |
| *CCL8* | 6355 |
| *CCND3* | 896 |
| *CCR1* | 1230 |
| *CCRL1* | 51554 |
| *CD274* | 29126 |
| *CD38* | 952 |
| *CD68* | 968 |
| *CD69* | 969 |
| *CD74* | 972 |
| *CD9* | 928 |
| *CDK17* | 5128 |
| *CDK18* | 5129 |
| *CDKN1A* | 1026 |
| *CEACAM1* | 634 |
| *CES1* | 1066 |
| *CFB* | 629 |
| *CHMP5* | 51510 |
| *CHORDC1* | 26973 |
| *CLEC2B* | 9976 |
| *CLEC4D* | 338339 |
| *CLEC4E* | 26253 |
| *CLEC7A* | 64581 |
| *CMAHP* | 8418 |
| *CNP* | 1267 |
| *CPT1A* | 1374 |

| | |
|---|---|
| *CRP* | 1401 |
| *CRY1* | 1407 |
| *CSRNP1* | 64651 |
| *CTCFL* | 140690 |
| *CTSS* | 1520 |
| *CX3CL1* | 6376 |
| *CXCL10* | 3627 |
| *CXCL11* | 6373 |
| *CXCL9* | 4283 |
| *CYP1B1* | 1545 |
| *CYP2J2* | 1573 |
| *DCP1A* | 55802 |
| *DDIT4* | 54541 |
| *DDX60* | 55601 |
| *DEFB1* | 1672 |
| *DNAH17* | 8632 |
| *DNAJA1* | 3301 |
| *DNAJC6* | 9829 |
| *DTX3L* | 151636 |
| *EHD4* | 30844 |
| *EIF3L* | 51386 |
| *ELF1* | 1997 |
| *ELOVL3* | 83401 |
| *EMR1* | 2015 |
| *EPAS1* | 2034 |
| *EPSTI1* | 94240 |
| *ERLIN1* | 10613 |
| *ETV6* | 2120 |
| *ETV7* | 51513 |
| *EXT1* | 2131 |
| *FAM122C* | 159091 |
| *FAM125B* | 89853 |
| *FAM134B* | 54463 |
| *FAM176A* | 84141 |
| *FAM46A* | 55603 |
| *FAM46C* | 54855 |
| *FAM70A* | 55026 |
| *FBXO6* | 26270 |
| *FCGR1A* | 2209 |
| *FFAR2* | 2867 |
| *FGD2* | 221472 |
| *FNDC3B* | 64778 |
| *FNDC4* | 64838 |
| *FSIP1* | 161835 |
| *FYB* | 2533 |
| *FZD5* | 7855 |
| *G6PC* | 2538 |
| *GALM* | 130589 |

| | |
|---|---|
| *GALNT2* | 2590 |
| *GBA3* | 57733 |
| *GBP1* | 2633 |
| *GBP2* | 2634 |
| *GBP3* | 2635 |
| *GBP5* | 115362 |
| *GCA* | 25801 |
| *GEM* | 2669 |
| *GIMAP2* | 26157 |
| *GJA4* | 2701 |
| *GK* | 2710 |
| *GLRX* | 2745 |
| *GMPR* | 2766 |
| *GNB4* | 59345 |
| *GSDMD* | 79792 |
| *GTPBP2* | 54676 |
| *GZMB* | 3002 |
| *HAPLN3* | 145864 |
| *HAS3* | 3038 |
| *HCP5* | 10866 |
| *HERC6* | 55008 |
| *HESX1* | 8820 |
| *HK2* | 3099 |
| *HLA-B* | 3106 |
| *HLA-C* | 3107 |
| *HLA-E* | 3133 |
| *HLA-F* | 3134 |
| *HLA-G* | 3135 |
| *HPSE* | 10855 |
| *HSP90AA1* | 3320 |
| *HSPA1A* | 3303 |
| *HSPA8* | 3312 |
| *HSPH1* | 10808 |
| *ID1* | 3397 |
| *ID2* | 3398 |
| *ID3* | 3399 |
| *IDO1* | 3620 |
| *IFI16* | 3428 |
| *IFI27* | 3429 |
| *IFI30* | 10437 |
| *IFI6* | 2537 |
| *IFIH1* | 64135 |
| *IFIT1* | 3434 |
| *IFIT2* | 3433 |
| *IFIT3* | 3437 |
| *IFIT5* | 24138 |
| *IFITM1* | 8519 |
| *IFITM2* | 10581 |

| | |
|---|---|
| *IFITM3* | 10410 |
| *IFNGR1* | 3459 |
| *IL11* | 3589 |
| *IL15* | 3600 |
| *IL1R1* | 3554 |
| *IL1RN* | 3557 |
| *IL22RA1* | 58985 |
| *IL4I1* | 259307 |
| *IL7* | 3574 |
| *IMPA2* | 3613 |
| *IRF2* | 3660 |
| *IRF9* | 10379 |
| *ISG15* | 9636 |
| *ISG20* | 3669 |
| *JUNB* | 3726 |
| *KBTBD8* | 84541 |
| *KIAA0040* | 9674 |
| *KIAA1644* | 85352 |
| *LAMP3* | 27074 |
| *LAP3* | 51056 |
| *LBP* | 3929 |
| *LGALS3* | 3958 |
| *LGALS9* | 3965 |
| *LGMN* | 5641 |
| *LIPA* | 3988 |
| *LMO2* | 4005 |
| *LNX1* | 84708 |
| *LOC152225* | 152225 |
| *LRG1* | 116844 |
| *LY6E* | 4061 |
| *LYAR* | 55646 |
| *LYSMD2* | 256586 |
| *MAB21L2* | 10586 |
| *MAK* | 4117 |
| *MAP2* | 4133 |
| *MAP3K14* | 9020 |
| *MAX* | 4149 |
| *MB21D1* | 115004 |
| *MCL1* | 4170 |
| *MKX* | 283078 |
| *MLKL* | 197259 |
| *MMP13* | 4322 |
| *MS4A4A* | 51338 |
| *MSR1* | 4481 |
| *MT1F* | 4494 |
| *MT1H* | 4496 |
| *MT1M* | 4499 |
| *MT1X* | 4501 |

| | |
|---|---|
| *MTHFD2L* | 441024 |
| *MX1* | 4599 |
| *MYD88* | 4615 |
| *NAMPT* | 10135 |
| *NAPA* | 8775 |
| *NAT8* | 9027 |
| *NCF1* | 653361 |
| *NCOA3* | 8202 |
| *NCOA7* | 135112 |
| *NDC80* | 10403 |
| *NEURL3* | 93082 |
| *NFIL3* | 4783 |
| *NID2* | 22795 |
| *NMI* | 9111 |
| *NRN1* | 51299 |
| *NT5C3* | 51251 |
| *NUP50* | 10762 |
| *NUPR1* | 26471 |
| *OAS1* | 4938 |
| *OAS2* | 4939 |
| *OASL* | 8638 |
| *OGFR* | 11054 |
| *OPTN* | 10133 |
| *OVOL1* | 5017 |
| *P2RY6* | 5031 |
| *PCGF5* | 84333 |
| *PDCD1LG2* | 80380 |
| *PDGFRL* | 5157 |
| *PDK1* | 5163 |
| *PFKFB3* | 5209 |
| *PHF11* | 51131 |
| *PHF15* | 23338 |
| *PIK3AP1* | 118788 |
| *PIM3* | 415116 |
| *PLAC8* | 51316 |
| *PLIN2* | 123 |
| *PLSCR1* | 5359 |
| *PLSCR4* | 57088 |
| *PMAIP1* | 5366 |
| *PML* | 5371 |
| *PMM2* | 5373 |
| *PPM1K* | 152926 |
| *PRAME* | 23532 |
| *PRICKLE4* | 29964 |
| *PRR15* | 222171 |
| *PSMB9* | 5698 |
| *PSME2* | 5721 |
| *PTMA* | 5757 |

| | |
|---|---|
| *PVRL4* | 81607 |
| *PXK* | 54899 |
| *RAB27A* | 5873 |
| *RAB39A* | 54734 |
| *RARRES3* | 5920 |
| *RASGEF1B* | 153020 |
| *RASGRP3* | 25780 |
| *RASSF4* | 83937 |
| *RBCK1* | 10616 |
| *RBM11* | 54033 |
| *REC8* | 9985 |
| *RET* | 5979 |
| *RGS1* | 5996 |
| *RGS22* | 26166 |
| *RIMS2* | 9699 |
| *RIPK2* | 8767 |
| *RNASE4* | 6038 |
| *RNF114* | 55905 |
| *RNF144A* | 9781 |
| *RNF19B* | 127544 |
| *RNF213* | 57674 |
| *RNF24* | 11237 |
| *RSAD2* | 91543 |
| *RTKN2* | 219790 |
| *RTP4* | 64108 |
| *RUFY4* | 285180 |
| *S100A8* | 6279 |
| *SAA1* | 6288 |
| *SAMD4A* | 23034 |
| *SAMHD1* | 25939 |
| *SCARB2* | 950 |
| *SCLT1* | 132320 |
| *SCO2* | 9997 |
| *SEC16B* | 89866 |
| *SECTM1* | 6398 |
| *SERPINB9* | 5272 |
| *SERPINE1* | 5054 |
| *SERPING1* | 710 |
| *SIRPA* | 140885 |
| *SLC16A4* | 9122 |
| *SLC1A1* | 6505 |
| *SLC25A30* | 253512 |
| *SLC2A12* | 154091 |
| *SLFN5* | 162394 |
| *SMAD3* | 4088 |
| *SNAI1* | 6615 |
| *SNN* | 8303 |
| *SOCS2* | 8835 |

| | |
|---|---|
| *SP100* | 6672 |
| *SP110* | 3431 |
| *SPATA13* | 221178 |
| *SPATS2L* | 26010 |
| *SPINK6* | 404203 |
| *SPSB1* | 80176 |
| *SPTLC2* | 9517 |
| *SQLE* | 6713 |
| *SSBP3* | 23648 |
| *ST3GAL4* | 6484 |
| *ST8SIA4* | 7903 |
| *STAP1* | 26228 |
| *STAT1* | 6772 |
| *STAT2* | 6773 |
| *STAT3* | 6774 |
| *STAT4* | 6775 |
| *SUN2* | 25777 |
| *SUSD3* | 203328 |
| *SUSD4* | 55061 |
| *TAGAP* | 117289 |
| *TAP1* | 6890 |
| *TAP2* | 6891 |
| *TCF4* | 6925 |
| *TFRC* | 7037 |
| *TGM2* | 7052 |
| *TIMP1* | 7076 |
| *TIMP2* | 7077 |
| *TLK2* | 11011 |
| *TLR2* | 7097 |
| *TLR3* | 7098 |
| *TLR7* | 51284 |
| *TMEM106A* | 113277 |
| *TMEM140* | 55281 |
| *TMEM171* | 134285 |
| *TMEM27* | 57393 |
| *TMEM51* | 55092 |
| *TMEM62* | 80021 |
| *TNFAIP3* | 7128 |
| *TNFAIP6* | 7130 |
| *TNFRSF10A* | 8797 |
| *TNFRSF1B* | 7133 |
| *TNFSF10* | 8743 |
| *TNFSF13* | 8741 |
| *TNFSF13B* | 10673 |
| *TOR1B* | 27348 |
| *TRAFD1* | 10906 |
| *TREX1* | 11277 |
| *TRIM21* | 6737 |

| | |
|---|---|
| *TRIM5* | 85363 |
| *TRIM56* | 81844 |
| *TRIM69* | 140691 |
| *TSPAN33* | 340348 |
| *TTC39B* | 158219 |
| *TYMP* | 1890 |
| *UBA7* | 7318 |
| *UBD* | 10537 |
| *UBE2L6* | 9246 |
| *ULK4* | 54986 |
| *UPP2* | 151531 |
| *USP18* | 11274 |
| *VAMP5* | 10791 |
| *VEGFC* | 7424 |
| *VMP1* | 81671 |
| *WARS* | 7453 |
| *ZBP1* | 81030 |
| *ZFP82* | 284406 |
| *ZNF385B* | 151126 |

identified based on experimental RNAseq (A549 or HTBE) and/or published datasets (Schogg

ins et al., 2011)

**Table S1B | Available ISGs**

cDNAs that were available as validated, full-sequence length clones (ORFeome library), from ISGs identified based on experimental RNAseq (A549 or HTBE) and/or published datasets (Schoggins et al., 2011)

| Gene Symbol | Gene ID |
|---|---|
| *ABLIM3* | 22885 |
| *ACSL5* | 51703 |
| *ACY3* | 91703 |
| *AHNAK2* | 113146 |
| *AIM2* | 9447 |
| *AKT3* | 10000 |
| *ALDH1A1* | 216 |
| *AMPH* | 273 |
| *ANGPTL1* | 9068 |
| *ANGPTL4* | 51129 |
| *ANKRD22* | 118932 |
| *APOBEC3A* | 200315 |
| *APOBEC3D* | 140564 |
| *APOBEC3F* | 200316 |
| *APOBEC3G* | 60489 |
| *APOL1* | 8542 |
| *APOL2* | 23780 |
| *APOL4* | 80832 |
| *APOL6* | 80830 |
| *AQP9* | 366 |
| *AREG* | 374 |
| *ARG2* | 384 |
| *ARHGAP17* | 55114 |
| *ARHGEF3* | 50650 |
| *ARNTL* | 406 |
| *ARRDC3* | 57561 |
| *ATF3* | 467 |
| *ATP10D* | 57205 |
| *AURKB* | 9212 |
| *B2M* | 567 |
| *B3GALNT1* | 8706 |
| *B4GALT5* | 9334 |
| *BAG1* | 573 |
| *BATF2* | 116071 |
| *BCL2L14* | 79370 |
| *BLNK* | 29760 |
| *BLVRA* | 644 |
| *BLZF1* | 8548 |
| *BMP4* | 652 |
| *BST2* | 684 |
| *BTC* | 685 |
| *BTN3A1* | 11119 |
| *BTN3A2* | 11118 |

| | |
|---|---|
| *BTN3A3* | 10384 |
| *C15orf48* | 84419 |
| *C17orf67* | 339210 |
| *C19orf66* | 55337 |
| *C1orf38* | 9473 |
| *C1R* | 715 |
| *C22orf28* | 51493 |
| *C3AR1* | 719 |
| *C3orf39* | 84892 |
| *C4orf33* | 132321 |
| *C5orf56* | 441108 |
| *C6orf141* | 135398 |
| *C9orf91* | 203197 |
| *CAPNS2* | 84290 |
| *CASP1* | 834 |
| *CASP10* | 843 |
| *CASP7* | 840 |
| *CBR3* | 874 |
| *CCDC109B* | 55013 |
| *CCDC75* | 253635 |
| *CCDC92* | 80212 |
| *CCL19* | 6363 |
| *CCL2* | 6347 |
| *CCL4* | 6351 |
| *CCL5* | 6352 |
| *CCL8* | 6355 |
| *CCND3* | 896 |
| *CCR1* | 1230 |
| *CCRL1* | 51554 |
| *CD274* | 29126 |
| *CD38* | 952 |
| *CD68* | 968 |
| *CD69* | 969 |
| *CD74* | 972 |
| *CD9* | 928 |
| *CDK17* | 5128 |
| *CDK18* | 5129 |
| *CDKN1A* | 1026 |
| *CEACAM1* | 634 |
| *CES1* | 1066 |
| *CFB* | 629 |
| *CHMP5* | 51510 |
| *CHORDC1* | 26973 |
| *CLEC2B* | 9976 |
| *CLEC4D* | 338339 |
| *CLEC4E* | 26253 |
| *CLEC7A* | 64581 |
| *CMAHP* | 8418 |

| | |
|---|---|
| *CNP* | 1267 |
| *CPT1A* | 1374 |
| *CRP* | 1401 |
| *CRY1* | 1407 |
| *CSRNP1* | 64651 |
| *CTCFL* | 140690 |
| *CTSS* | 1520 |
| *CX3CL1* | 6376 |
| *CXCL10* | 3627 |
| *CXCL11* | 6373 |
| *CXCL9* | 4283 |
| *CYP1B1* | 1545 |
| *CYP2J2* | 1573 |
| *DCP1A* | 55802 |
| *DDIT4* | 54541 |
| *DDX60* | 55601 |
| *DEFB1* | 1672 |
| *DNAH17* | 8632 |
| *DNAJA1* | 3301 |
| *DNAJC6* | 9829 |
| *DTX3L* | 151636 |
| *EHD4* | 30844 |
| *EIF3L* | 51386 |
| *ELF1* | 1997 |
| *ELOVL3* | 83401 |
| *EMR1* | 2015 |
| *EPAS1* | 2034 |
| *EPSTI1* | 94240 |
| *ERLIN1* | 10613 |
| *ETV6* | 2120 |
| *ETV7* | 51513 |
| *EXT1* | 2131 |
| *FAM122C* | 159091 |
| *FAM125B* | 89853 |
| *FAM134B* | 54463 |
| *FAM176A* | 84141 |
| *FAM46A* | 55603 |
| *FAM46C* | 54855 |
| *FAM70A* | 55026 |
| *FBXO6* | 26270 |
| *FCGR1A* | 2209 |
| *FFAR2* | 2867 |
| *FGD2* | 221472 |
| *FNDC3B* | 64778 |
| *FNDC4* | 64838 |
| *FSIP1* | 161835 |
| *FYB* | 2533 |
| *FZD5* | 7855 |

| | |
|---|---|
| *G6PC* | 2538 |
| *GALM* | 130589 |
| *GALNT2* | 2590 |
| *GBA3* | 57733 |
| *GBP1* | 2633 |
| *GBP2* | 2634 |
| *GBP3* | 2635 |
| *GBP5* | 115362 |
| *GCA* | 25801 |
| *GEM* | 2669 |
| *GIMAP2* | 26157 |
| *GJA4* | 2701 |
| *GK* | 2710 |
| *GLRX* | 2745 |
| *GMPR* | 2766 |
| *GNB4* | 59345 |
| *GSDMD* | 79792 |
| *GTPBP2* | 54676 |
| *GZMB* | 3002 |
| *HAPLN3* | 145864 |
| *HAS3* | 3038 |
| *HCP5* | 10866 |
| *HERC6* | 55008 |
| *HESX1* | 8820 |
| *HK2* | 3099 |
| *HLA-B* | 3106 |
| *HLA-C* | 3107 |
| *HLA-E* | 3133 |
| *HLA-F* | 3134 |
| *HLA-G* | 3135 |
| *HPSE* | 10855 |
| *HSP90AA1* | 3320 |
| *HSPA1A* | 3303 |
| *HSPA8* | 3312 |
| *HSPH1* | 10808 |
| *ID1* | 3397 |
| *ID2* | 3398 |
| *ID3* | 3399 |
| *IDO1* | 3620 |
| *IFI16* | 3428 |
| *IFI27* | 3429 |
| *IFI30* | 10437 |
| *IFI6* | 2537 |
| *IFIH1* | 64135 |
| *IFIT1* | 3434 |
| *IFIT2* | 3433 |
| *IFIT3* | 3437 |
| *IFIT5* | 24138 |

| | |
|---|---|
| *IFITM1* | 8519 |
| *IFITM2* | 10581 |
| *IFITM3* | 10410 |
| *IFNGR1* | 3459 |
| *IL11* | 3589 |
| *IL15* | 3600 |
| *IL1R1* | 3554 |
| *IL1RN* | 3557 |
| *IL22RA1* | 58985 |
| *IL4I1* | 259307 |
| *IL7* | 3574 |
| *IMPA2* | 3613 |
| *IRF2* | 3660 |
| *IRF9* | 10379 |
| *ISG15* | 9636 |
| *ISG20* | 3669 |
| *JUNB* | 3726 |
| *KBTBD8* | 84541 |
| *KIAA0040* | 9674 |
| *KIAA1644* | 85352 |
| *LAMP3* | 27074 |
| *LAP3* | 51056 |
| *LBP* | 3929 |
| *LGALS3* | 3958 |
| *LGALS9* | 3965 |
| *LGMN* | 5641 |
| *LIPA* | 3988 |
| *LMO2* | 4005 |
| *LNX1* | 84708 |
| *LOC152225* | 152225 |
| *LRG1* | 116844 |
| *LY6E* | 4061 |
| *LYAR* | 55646 |
| *LYSMD2* | 256586 |
| *MAB21L2* | 10586 |
| *MAK* | 4117 |
| *MAP2* | 4133 |
| *MAP3K14* | 9020 |
| *MAX* | 4149 |
| *MB21D1* | 115004 |
| *MCL1* | 4170 |
| *MKX* | 283078 |
| *MLKL* | 197259 |
| *MMP13* | 4322 |
| *MS4A4A* | 51338 |
| *MSR1* | 4481 |
| *MT1F* | 4494 |
| *MT1H* | 4496 |

| | |
|---|---|
| *MT1M* | 4499 |
| *MT1X* | 4501 |
| *MTHFD2L* | 441024 |
| *MX1* | 4599 |
| *MYD88* | 4615 |
| *NAMPT* | 10135 |
| *NAPA* | 8775 |
| *NAT8* | 9027 |
| *NCF1* | 653361 |
| *NCOA3* | 8202 |
| *NCOA7* | 135112 |
| *NDC80* | 10403 |
| *NEURL3* | 93082 |
| *NFIL3* | 4783 |
| *NID2* | 22795 |
| *NMI* | 9111 |
| *NRN1* | 51299 |
| *NT5C3* | 51251 |
| *NUP50* | 10762 |
| *NUPR1* | 26471 |
| *OAS1* | 4938 |
| *OAS2* | 4939 |
| *OASL* | 8638 |
| *OGFR* | 11054 |
| *OPTN* | 10133 |
| *OVOL1* | 5017 |
| *P2RY6* | 5031 |
| *PCGF5* | 84333 |
| *PDCD1LG2* | 80380 |
| *PDGFRL* | 5157 |
| *PDK1* | 5163 |
| *PFKFB3* | 5209 |
| *PHF11* | 51131 |
| *PHF15* | 23338 |
| *PIK3AP1* | 118788 |
| *PIM3* | 415116 |
| *PLAC8* | 51316 |
| *PLIN2* | 123 |
| *PLSCR1* | 5359 |
| *PLSCR4* | 57088 |
| *PMAIP1* | 5366 |
| *PML* | 5371 |
| *PMM2* | 5373 |
| *PPM1K* | 152926 |
| *PRAME* | 23532 |
| *PRICKLE4* | 29964 |
| *PRR15* | 222171 |
| *PSMB9* | 5698 |

| | |
|---|---|
| *PSME2* | 5721 |
| *PTMA* | 5757 |
| *PVRL4* | 81607 |
| *PXK* | 54899 |
| *RAB27A* | 5873 |
| *RAB39A* | 54734 |
| *RARRES3* | 5920 |
| *RASGEF1B* | 153020 |
| *RASGRP3* | 25780 |
| *RASSF4* | 83937 |
| *RBCK1* | 10616 |
| *RBM11* | 54033 |
| *REC8* | 9985 |
| *RET* | 5979 |
| *RGS1* | 5996 |
| *RGS22* | 26166 |
| *RIMS2* | 9699 |
| *RIPK2* | 8767 |
| *RNASE4* | 6038 |
| *RNF114* | 55905 |
| *RNF144A* | 9781 |
| *RNF19B* | 127544 |
| *RNF213* | 57674 |
| *RNF24* | 11237 |
| *RSAD2* | 91543 |
| *RTKN2* | 219790 |
| *RTP4* | 64108 |
| *RUFY4* | 285180 |
| *S100A8* | 6279 |
| *SAA1* | 6288 |
| *SAMD4A* | 23034 |
| *SAMHD1* | 25939 |
| *SCARB2* | 950 |
| *SCLT1* | 132320 |
| *SCO2* | 9997 |
| *SEC16B* | 89866 |
| *SECTM1* | 6398 |
| *SERPINB9* | 5272 |
| *SERPINE1* | 5054 |
| *SERPING1* | 710 |
| *SIRPA* | 140885 |
| *SLC16A4* | 9122 |
| *SLC1A1* | 6505 |
| *SLC25A30* | 253512 |
| *SLC2A12* | 154091 |
| *SLFN5* | 162394 |
| *SMAD3* | 4088 |
| *SNAI1* | 6615 |

| | |
|---|---|
| *SNN* | 8303 |
| *SOCS2* | 8835 |
| *SP100* | 6672 |
| *SP110* | 3431 |
| *SPATA13* | 221178 |
| *SPATS2L* | 26010 |
| *SPINK6* | 404203 |
| *SPSB1* | 80176 |
| *SPTLC2* | 9517 |
| *SQLE* | 6713 |
| *SSBP3* | 23648 |
| *ST3GAL4* | 6484 |
| *ST8SIA4* | 7903 |
| *STAP1* | 26228 |
| *STAT1* | 6772 |
| *STAT2* | 6773 |
| *STAT3* | 6774 |
| *STAT4* | 6775 |
| *SUN2* | 25777 |
| *SUSD3* | 203328 |
| *SUSD4* | 55061 |
| *TAGAP* | 117289 |
| *TAP1* | 6890 |
| *TAP2* | 6891 |
| *TCF4* | 6925 |
| *TFRC* | 7037 |
| *TGM2* | 7052 |
| *TIMP1* | 7076 |
| *TIMP2* | 7077 |
| *TLK2* | 11011 |
| *TLR2* | 7097 |
| *TLR3* | 7098 |
| *TLR7* | 51284 |
| *TMEM106A* | 113277 |
| *TMEM140* | 55281 |
| *TMEM171* | 134285 |
| *TMEM27* | 57393 |
| *TMEM51* | 55092 |
| *TMEM62* | 80021 |
| *TNFAIP3* | 7128 |
| *TNFAIP6* | 7130 |
| *TNFRSF10A* | 8797 |
| *TNFRSF1B* | 7133 |
| *TNFSF10* | 8743 |
| *TNFSF13* | 8741 |
| *TNFSF13B* | 10673 |
| *TOR1B* | 27348 |
| *TRAFD1* | 10906 |

| | |
|---|---|
| *TREX1* | 11277 |
| *TRIM21* | 6737 |
| *TRIM5* | 85363 |
| *TRIM56* | 81844 |
| *TRIM69* | 140691 |
| *TSPAN33* | 340348 |
| *TTC39B* | 158219 |
| *TYMP* | 1890 |
| *UBA7* | 7318 |
| *UBD* | 10537 |
| *UBE2L6* | 9246 |
| *ULK4* | 54986 |
| *UPP2* | 151531 |
| *USP18* | 11274 |
| *VAMP5* | 10791 |
| *VEGFC* | 7424 |
| *VMP1* | 81671 |
| *WARS* | 7453 |
| *ZBP1* | 81030 |
| *ZFP82* | 284406 |
| *ZNF385B* | 151126 |

**Table S2 | Overexpression Screen Hits**

293Tcells were transfected with each of the indicated ISGs along with ACE2 and TMPRSS2. At 36 h post-transfection were infected with SARS-CoV-2 (MOI = 0.0625) for 40 h prior to immunostaining for viral N. Column E shows mean infectivity (% N+ positive cells) relative to negative control CAT.

| Gene Symbol | gene ID | Description | Normalized infection |
|---|---|---|---|
| *ANGPTL4* | 51129 | angiopoietin like 4 | 0.464 |
| *APOL2* | 23780 | apolipoprotein L2 | 0.598 |
| *APOL4* | 80832 | apolipoprotein L4 | 0.250 |
| *ARNTL* | 406 | aryl hydrocarbon receptor nuclear translocator like | 0.489 |
| *B4GALT5* | 9334 | beta-1,4-galactosyltransferase 5 | 0.425 |
| *BST2* | 684 | bone marrow stromal cell antigen 2 | 0.065 |
| *C9orf91* | 203197 | transmembrane protein 268 | 0.515 |
| *CASP7* | 840 | caspase 7 | 0.598 |
| *CCND3* | 896 | cyclin D3 | 0.012 |
| *CLEC4D* | 338339 | C-type lectin domain family 4 member D | 0.718 |
| *CNP* | 1267 | 2',3'-cyclic nucleotide 3' phosphodiesterase | 0.668 |
| *CRP* | 1401 | C-reactive protein | 0.462 |
| *DDX60* | 55601 | DExD/H-box helicase 60 | 0.603 |
| *DNAJC6* | 9829 | DnaJ heat shock protein family (Hsp40) member C6 | 0.646 |
| *ELF1* | 1997 | E74 like ETS transcription factor 1 | 0.295 |
| *ERLIN1* | 10613 | ER lipid raft associated 1 | 0.559 |
| *ETV6* | 2120 | ETS variant transcription factor 6 | 0.638 |
| *FAM134B* | 54463 | reticulophagy regulator 1 | 0.508 |
| *FAM46A* | 55603 | terminal nucleotidyltransferase 5A | 0.644 |
| *FAM46C* | 54855 | terminal nucleotidyltransferase 5C | 0.655 |
| *FGD2* | 221472 | FYVE, RhoGEF and PH domain containing 2 | 0.493 |
| *FNDC4* | 64838 | fibronectin type III domain containing 4 | 0.666 |
| *FZD5* | 7855 | frizzled class receptor 5 | 0.552 |
| *GBA3* | 57733 | glucosylceramidase beta 3 (gene/pseudogene) | 0.650 |
| *GBP3* | 2635 | guanylate binding protein 3 | 0.273 |
| *GNB4* | 59345 | G protein subunit beta 4 | 0.450 |
| *GSDMD* | 79792 | gasdermin D | 0.766 |
| *HSPA8* | 3312 | heat shock protein family A (Hsp70) member 8 | 0.154 |
| *IFIT1* | 3434 | IFN-induced protein with tetratricopeptide repeats 1 | 0.488 |
| *IFIT3* | 3437 | IFN-induced protein with tetratricopeptide repeats 3 | 0.597 |
| *IFIT5* | 24138 | IFN-induced protein with tetratricopeptide repeats 5 | 0.569 |
| *IFITM2* | 10581 | interferon induced transmembrane protein 2 | 0.276 |
| *IFITM3* | 10410 | interferon induced transmembrane protein 3 | 0.634 |
| *IL11* | 3589 | interleukin 11 | 0.317 |
| *IL4I1* | 259307 | interleukin 4 induced 1 | 0.404 |
| *ISG15* | 9636 | ISG15 ubiquitin like modifier | 0.426 |
| *ISG20* | 3669 | interferon stimulated exonuclease gene 20 | 0.503 |
| *LOC152225* | 152225 | long intergenic non-protein coding RNA 2085 | 0.551 |
| *LY6E* | 4061 | lymphocyte antigen 6 family member E | 0.242 |
| *MAX* | 4149 | MYC associated factor X | 0.676 |
| *MLKL* | 197259 | mixed lineage kinase domain like pseudokinase | 0.064 |
| *MSR1* | 4481 | macrophage scavenger receptor 1 | 0.241 |

| | | | |
|---|---|---|---|
| ***MYD88*** | 4615 | MYD88 innate immune signal transduction adaptor | 0.621 |
| ***NAPA*** | 8775 | NSF attachment protein alpha | 0.256 |
| ***NRN1*** | 51299 | neuritin 1 | 0.653 |
| ***NT5C3*** | 51251 | 5'-nucleotidase, cytosolic IIIA | 0.476 |
| ***PHF15*** | 23338 | jade family PHD finger 2 | 0.562 |
| ***PIK3AP1*** | 118788 | phosphoinositide-3-kinase adaptor protein 1 | 0.614 |
| ***RAB27A*** | 5873 | RAB27A, member RAS oncogene family | 0.574 |
| ***RAB39A*** | 54734 | RAB39A, member RAS oncogene family | 0.100 |
| ***REC8*** | 9985 | REC8 meiotic recombination protein | 0.604 |
| ***RGS22*** | 26166 | regulator of G protein signaling 22 | 0.697 |
| ***RSAD2*** | 91543 | radical S-adenosyl methionine domain containing 2 | 0.596 |
| ***SPATA13*** | 221178 | spermatogenesis associated 13 | 0.646 |
| ***SPATS2L*** | 26010 | spermatogenesis associated serine rich 2 like | 0.538 |
| ***ST3GAL4*** | 6484 | ST3 beta-galactoside alpha-2,3-sialyltransferase 4 | 0.456 |
| ***STAT1*** | 6772 | signal transducer and activator of transcription 1 | 0.572 |
| ***STAT2*** | 6773 | signal transducer and activator of transcription 2 | 0.578 |
| ***SUSD3*** | 203328 | sushi domain containing 3 | 0.482 |
| ***TAGAP*** | 117289 | T cell activation RhoGTPase activating protein | 0.591 |
| ***TRIM21*** | 6737 | tripartite motif containing 21 | 0.665 |
| ***UBD*** | 10537 | ubiquitin D | 0.583 |
| ***UPP2*** | 151531 | uridine phosphorylase 2 | 0.683 |
| ***USP18*** | 11274 | ubiquitin specific peptidase 18 | 0.436 |
| ***ZBP1*** | 81030 | Z-DNA binding protein 1 | 0.352 |

**Table S3 | Lentivirus validated hits**

293T-ACE2 stably expressing each of the indicated ISGs were infected with SARS-CoV-2 (MOI = 0.25) for 40 h prior to immunostaining for SARS-CoV-2 N protein. Column C shows average relative infectivity (% N+ positive cells) to parental cells across 3 independent experiments. Statistical significance (p value, column D) was calculated using one-way ANOVA with Sidak´s multiple comparison post-hoc test

| Gene Symbol | Gene ID | Normalized infection | p value |
|---|---|---|---|
| *APOL2* | 23780 | 0.246 | 0.0012 |
| *B4GALT5* | 9334 | 0.270 | 0.0011 |
| *BST2* | 684 | 0.083 | <0.0001 |
| *C9orf91* | 203197 | 0.380 | 0.0231 |
| *CLEC4D* | 338339 | 0.016 | <0.0001 |
| *CNP* | 1267 | 0.185 | <0.0001 |
| *DNAJC6* | 9829 | 0.099 | <0.0001 |
| *ELF1* | 1997 | 0.059 | <0.0001 |
| *ERLIN1* | 10613 | 0.167 | <0.0001 |
| *ETV6* | 2120 | 0.288 | 0.0058 |
| *FAM46A* | 55603 | 0.045 | <0.0001 |
| *FAM46C* | 54855 | 0.069 | <0.0001 |
| *FZD5* | 7855 | 0.404 | 0.0328 |
| *GNB4* | 59345 | 0.170 | <0.0001 |
| *HSPA8* | 3312 | 0.147 | <0.0001 |
| *IFIT3* | 3437 | 0.108 | <0.0001 |
| *IFIT5* | 24138 | 0.365 | 0.0233 |
| *IFITM2* | 10581 | 0.233 | 0.0006 |
| *IFITM3* | 10410 | 0.343 | 0.0285 |
| *ISG15* | 9636 | 0.347 | 0.023 |
| *ISG20* | 3669 | 0.421 | 0.0589 |
| *LOC152225* | 152225 | 0.157 | <0.0001 |
| *LY6E* | 4061 | 0.232 | <0.0001 |
| *MSR1* | 4481 | 0.161 | <0.0001 |
| *NRN1* | 51299 | 0.464 | 0.0493 |
| *NT5C3* | 51251 | 0.211 | <0.0001 |
| *PHF15* | 23338 | 0.419 | 0.0589 |
| *RAB27A* | 5873 | 0.131 | <0.0001 |
| *REC8* | 9985 | 0.039 | <0.0001 |
| *RGS22* | 26166 | 0.075 | <0.0001 |
| *SPATA13* | 221178 | 0.372 | 0.0328 |
| *SPATS2L* | 26010 | 0.070 | <0.0001 |
| *STAT1* | 6772 | 0.247 | 0.0011 |
| *TAGAP* | 117289 | 0.196 | <0.0001 |
| *TRIM21* | 6737 | 0.135 | <0.0001 |
| *UBD* | 10537 | 0.371 | 0.0562 |
| *ZBP1* | 81030 | 0.317 | 0.0138 |
